# Multiscale Cloud-Based Pipeline for Neuronal Electrophysiology Analysis and Visualization

**DOI:** 10.1101/2024.11.14.623530

**Authors:** Jinghui Geng, Kateryna Voitiuk, David F. Parks, Ash Robbins, Alex Spaeth, Jessica L. Sevetson, Sebastian Hernandez, Hunter E. Schweiger, John P. Andrews, Spencer T. Seiler, Matthew A.T. Elliott, Edward F. Chang, Tomasz J. Nowakowski, Rob Currie, Mohammed A. Mostajo-Radji, David Haussler, Tal Sharf, Sofie R. Salama, Mircea Teodorescu

**Affiliations:** Department of Electrical and Computer Engineering, University of California Santa Cruz, Santa Cruz, CA 95064, USA; Department of Biomolecular Engineering, University of California Santa Cruz, Santa Cruz, CA 95064, USA; Department of Molecular, Cell, and Developmental Biology, University of California Santa Cruz, Santa Cruz, CA 95064, USA; Genomics Institute, University of California Santa Cruz, Santa Cruz, CA 95064, USA; Department of Neurological Surgery, University of California San Francisco, San Francisco, CA 94143, USA; Weill Institute for Neurosciences, University of California San Francisco, San Francisco, CA 94143, USA; The Eli and Edythe Broad Center of Regeneration Medicine and Stem Cell Research, University of California San Francisco, San Francisco, CA 94143, USA; Department of Psychiatry and Behavioral Sciences, University of California San Francisco, San Francisco, CA 94143, USA; Department of Anatomy, University of California San Francisco, San Francisco, CA 94143, USA

**Keywords:** microelectrode arrays, electrophysiology recording analysis, data pipeline, Internet of Things, cloud computing, containerization, neuronal cultures

## Abstract

Electrophysiology offers a high-resolution method for real-time measurement of neural activity. Longitudinal recordings from high-density microelectrode arrays (HD-MEAs) can be of considerable size for local storage and of substantial complexity for extracting neural features and network dynamics. Analysis is often demanding due to the need for multiple software tools with different runtime dependencies. To address these challenges, we developed an open-source cloud-based pipeline to store, analyze, and visualize neuronal electrophysiology recordings from HD-MEAs. This pipeline is dependency agnostic by utilizing cloud storage, cloud computing resources, and an Internet of Things messaging protocol. We containerized the services and algorithms to serve as scalable and flexible building blocks within the pipeline. In this paper, we applied this pipeline on two types of cultures, cortical organoids and *ex vivo* brain slice recordings to show that this pipeline simplifies the data analysis process and facilitates understanding neuronal activity.

## INTRODUCTION

Recent advances in hardware and software platforms for neuronal recordings have enabled simultaneous recording of neuronal activity with high spatial and temporal resolution across various samples, including brain slices ^1,2^, 2D cultures ^3–5^, and 3D cerebral organoids ^6,7^. These technologies facilitate comprehensive studies of brain function, neurodevelopment, and network topology ^8–10^. However, the exponential growth in data volume and complexity ^11–13^ presents significant challenges in data storage, processing, and analysis. Recordings, images, and analysis results can consume substantial storage on computers and hard drives. Interpreting this multi-dimensional data requires specialized algorithms and tools to extract single neuronal unit activity, visualize firing patterns, and understand neuronal network-level information ^14–16^. While efforts have been made to unify standards in electrophysiology, biologists still face difficulties performing comprehensive analyses.

Spike sorting algorithms are crucial for analyzing multi-electrode array (MEA) recordings ^17–21^, identifying and categorizing individual neuronal spikes from raw voltage traces to analyze neuronal features ^22–25^ and network dynamics ^26,27^. While various software tools have been developed to process MEA recordings and visualize neuronal features ^23,28–33^, challenges persist due to differing programming languages, limited user support, and compatibility issues. Although integrated platforms offer end-to-end analysis capabilities, they may restrict custom data manipulation, requiring researchers to develop their own workflows and navigate steep learning curves for effective data interpretation.

Cloud computing enables processing a large amount of data in parallel by utilizing abundant resources while still being a cost-effective solution ^34–36^. Cloud-based storage can address the issue of massive experimental data filling up local disks. It also provides extensive data sharing ability for collaborations across research labs. Infrastructures and web platforms have been developed to store and analyze various types of data, including electrophysiology, neuroimaging, and sequencing ^37–42^. These platforms are designed to benefit the broader neuroscience community, emphasizing data publication and sharing ^43,44^. A research laboratory-oriented data platform is needed to support consistent experiments and data processing.

The Internet of Things (IoT) has made a significant impact in many fields, including healthcare ^45,46^ and, in recent years, has been applied to cellular biology ^47–49^ and *in vitro* electrophysiology experiments ^50–53^. Its resource efficiency enables the messaging protocol to work across different hardware, allowing networks to grow from a few devices to a large number without compromising performance.

We developed a cloud-based pipeline for electrophysiology data storage, processing, and sharing to facilitate the day-to-day research. We used containerization as the minimum building block. The IoT messaging services and data analysis algorithms are packaged into individual containers. The IoT services run on a web server to stream data, monitor processing tasks, and communicate with researchers through user interfaces. We applied Kubernetes ^54^ to orchestrate the analysis containers on the cloud computing clusters. By using cloud computing resources, the pipeline can process a large number of datasets with different algorithms in parallel, optimizing resource utilization, scalability, and flexibility. Moreover, we lower costs by replacing local computing hardware, such as CPUs and GPUs, with cloud-based technology. We also remove the barriers to data analysis by providing user interfaces, minimizing the software setup process, and making the Python code open source. The pipeline provides a suite of algorithms, including spike sorting, autocuration of putative neural units, visualization, and downstream analyses for specific goals using the curated data. We tested this pipeline with two applications. First, we analyzed mouse cortical organoid longitudinal recordings, 10 minutes long, one hour apart, over a 7-day period. This demonstrated the utility of our approach for neuron tracking. Second, we applied the pipeline to study optogenetic modulation of epileptiform activity in human hippocampus slices, contributing to our understanding and potential treatment of neurological diseases.

## RESULTS

Our platform allows users to upload recordings from electrophysiology devices directly to cloud storage. The data is organized by experiment date and is annotated with automatically extracted as well as user-specified metadata. The pipeline can be scaled up as algorithms and services are containerized, making it easy to integrate new analytical tools as they become available. The pipeline supports multiple data processing paradigms to accommodate diverse research requirements. The graphical interface allows users to initiate, monitor, and visualize data processing after upload, offering multimodal analysis and result downloads. An integrated IoT messaging service connects users, local recording devices, and the cloud, streamlining workflow.

### Framework Design

The pipeline is generic and capable of processing data from any electrophysiology platform that uses HDF5 and NWB ^55–57^ formats. In this paper, we tested it with data generated by a MaxOne HD-MEA (MaxWell Biosystems) ^58^. The system has 26,400 electrodes in a 2.10 x 3.85mm^2^ area. It supports data collection from 1,020 channels and can simulate 32 channels simultaneously at a 20 kHz sample rate. Together with a small inter-electrode pitch (17.5 *µ*m), the system provides high temporal and spatial resolution, where the activity of a typical neuron will be recorded on multiple pads. We utilize Ceph/S3^59^ and the National Research Platform (NRP) ^60,61^ computing clusters for data storage and processing.

The overview of the platform is shown in Figure 1. Neuronal tissue culture activity data is collected on a MaxWell MEA headstage, connected to a local computer running MaxLab software (Figure 1A). After recording, datasets are streamed to S3 and the data uploader generates corresponding metadata and maintains the applicable S3 file structure for these datasets (Figure 1B). Upon completion, an MQTT message is sent from the data uploader to the processing service – the job listener. This message contains the experiment identifiers and the image of the dockerized algorithm. The listener parses the message to gather the S3 paths for each dataset and calls the Kubernetes-Python API to deploy data processing jobs to the NRP computing cluster 15. The pipeline provides several containerized data processing applications, including spike sorting, data curation, and visualization. Once a job is completed on the NRP, the result is saved to S3 (Figure 1C). Researchers can access and download these results through the user interface.

**Figure 1:**
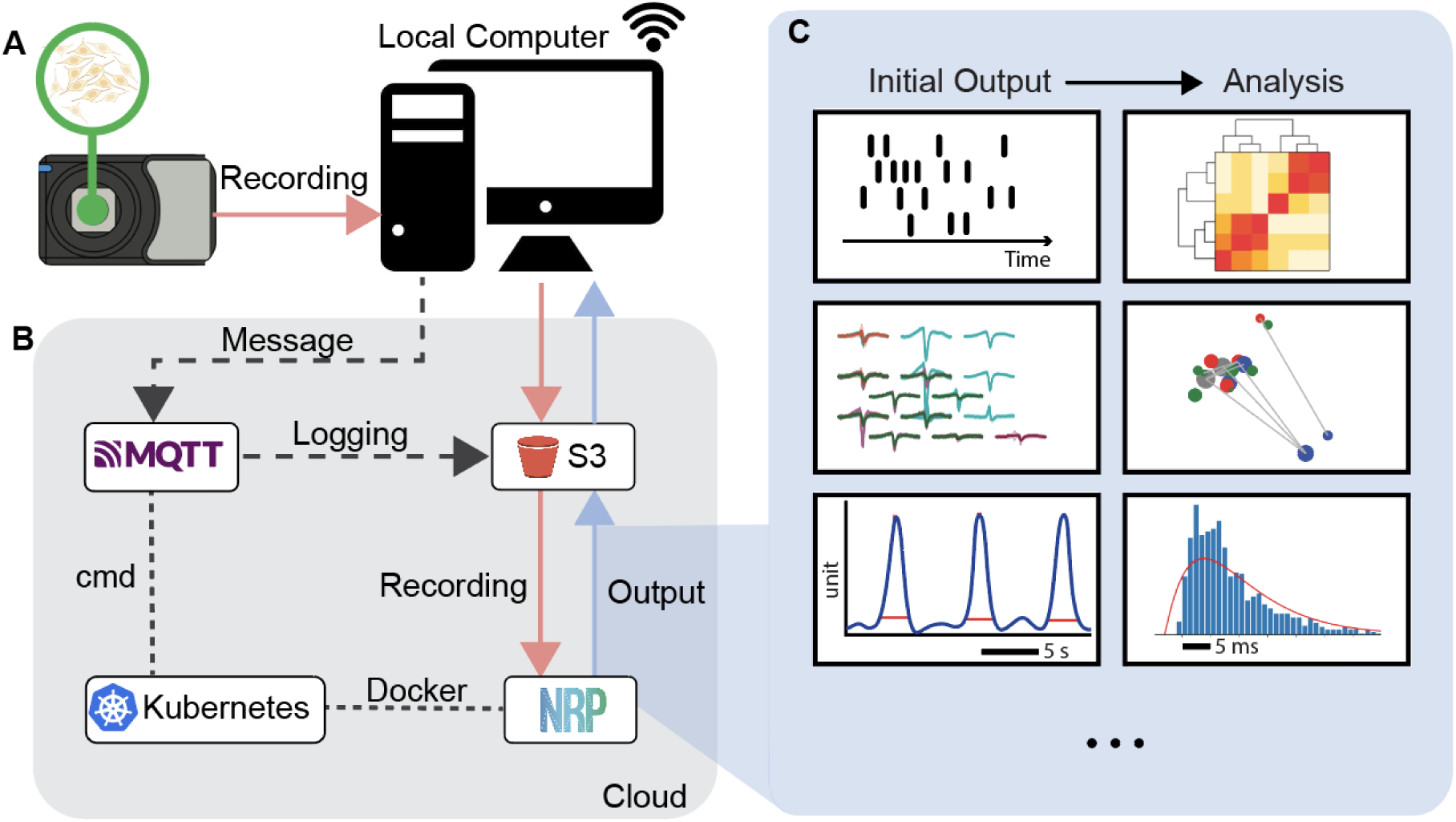
Cloud-based electrophysiology data processing pipeline architecture. (A) Electrophysiology data from neuronal cultures is recorded on a local computer. Different neuronal cultures and their recordings are shown in Figures 4 and 7. (B) Once the dataset is saved, it is uploaded to a uniquely identified data bucket S3 for permanent storage using the Uploader. An MQTT message is simultaneously sent to the job listener service to initiate data processing jobs. These jobs run containerized algorithms and are launched on the National Research Platform (NRP) computing cluster using Kubernetes. Results, including post-processed data and figures, are saved back to S3. (C) The analysis outputs various interactive analytical figures for each dataset’s network features and single-unit activity.

To make the pipeline accessible to non-programmers, we have developed user interfaces for managing and interacting with both local and remote data processes (Figure 2A). Through these interfaces, researchers can have complete control over their data while bridging the gap between running complex algorithms and requiring extensive programming knowledge or technical expertise.

**Figure 2:**
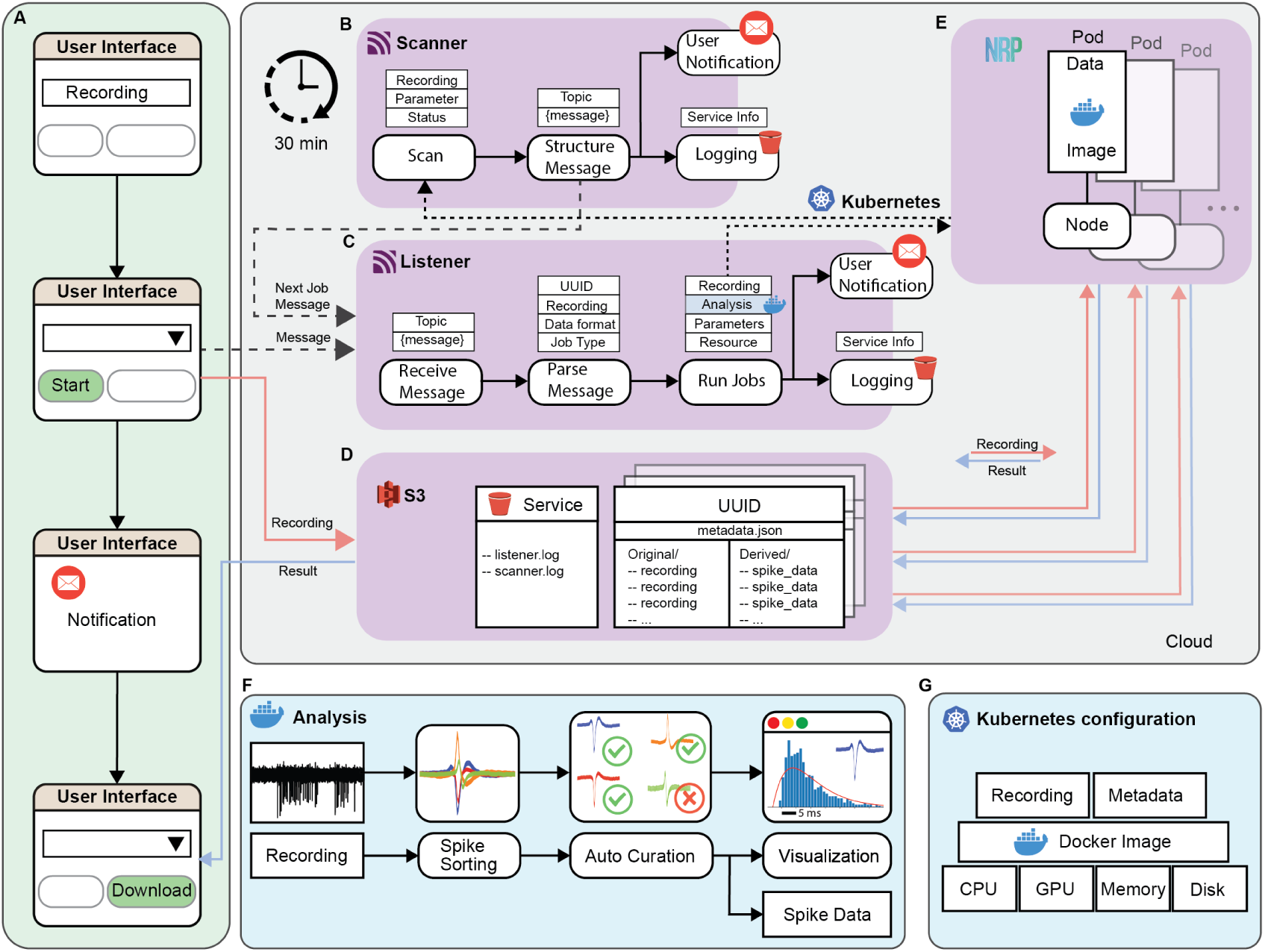
Pipeline components and workflow. (A) The user interface allows researchers to upload their electrophysiology recordings to cloud storage, initiate data processing jobs, receive notification upon completion, and download results to local computers. (B) MQTT-based job scanner service monitors job status on the NRP, sends a message to the listener for the next job, and notifies users. (C) MQTT-based job listener service that subscribes to specific topics to run data processing jobs. When the service receives a message, it parses the JSON format to extract experiment identifiers and computing requirements, then deploys jobs to NRP through Python-Kubernetes API. Both scanner and listener services update their status to S3 log files on a scheduled basis. (D) S3 file structure for service logging and experiment data. Log files are human-readable text files that track service status. Experiment data is stored in batches, each with a unique identifier (UUID), metadata file, “original” bucket for experiment data, and “derived” bucket for analysis outputs. (E) Computing cluster (NRP) for running containerized jobs using Kubernetes. (F) An analysis container for batch processing is capable of loading electrophysiology recordings, running spike sorting and autocuration algorithms, producing visualization figures, and generating Numpy files for single units. (G) Kubernetes configuration for job deployment to a computing cluster.

These interfaces include a data uploader, a Dashboard webpage, and a Slack channel, each serving distinct purposes while bridging local data collection, cloud-based data manipulation, and user notifications. The data uploader, installable on local laptops, enables users to upload electrophysiology recordings to S3 storage and initiate batch processing jobs with predefined parameters after the experiment is finished. The web Dashboard, accessible from any internet-connected device, provides access to existing S3 data for both batch and chained jobs. The data uploader and the Dashboard support downloading files from S3 to local directories. Additionally, the Dashboard features a visualization page displaying post-processing figures of selected recordings. A Slack channel is used to post status notifications for data processing jobs. Detailed descriptions of these user interfaces are provided in the Methods section. Screenshots of the applications are shown in Supplementary Figure S2, S3.

For cloud integration, we used the Message Queuing Telemetry Transport (MQTT) messaging protocol, a lightweight publish-subscribe protocol designed for Internet-of-Things (IoT) applications. This approach reduces the dependency requirements for edge devices to run cloud-computing jobs. A local computer can utilize the pipeline as long as it can run a Python environment and has a network connection. We have designed job listener and scanner services to run and monitor jobs on the cloud (Figure 2B,C).

For cloud computing, we used the National Research Platform (NRP), a distributed commodity compute cluster based on Kubernetes and the Ceph distributed file system. It has special CPUs and GPUs for data science, simulations, and machine learning. This setup allows for parallel data processing and can help reduce the computing infrastructure cost of individual labs.

We have a job scanner (Figure 2B) that checks on data processing jobs in the cloud every 30 minutes. It updates a list of current job statuses using the Kubernetes Python API. The scanner reads job names and information, which are named based on the dataset or a job list. This helps the scanner find the correct information in the NRP. The scanner then updates the listener and the user about how jobs are progressing.

To keep the flexibility of data processing, we implemented two types of jobs: batch processing and chained jobs. Chained jobs run through several steps on different data, with subsequent processing dependent on prior results. When the scanner detects a status change in a chained job, it sends a message to the listener to update the corresponding job look-up table and initiate the next processing step. Concurrently, it notifies the user about completing the prior job and the start of the next. This notification is done through the “slack-bridge” service. For completed batch processing jobs, the scanner sends only a user notification. After a message is sent, finished jobs are removed from the scanner’s memory to prevent duplicate notifications.

The job listener (Figure 2C) receives messages from both the user interface and the scanner. It also sends user notifications to the “ephys-pipeline” Slack channel. The primary function of the job listener is to initiate cloud computing jobs. Upon receiving a run job message, the listener parses it to extract the data path, data format, parameter setting, and job type (analysis algorithms). The listener then calls functions from a Python Kubernetes object (Figure 2) to allocate computing resources on NRP and the appropriate analysis docker image for each dataset. This object creates a job on the NRP and a pod within each job. Finally, it sends a “job created” notification to the Slack channel. Both the scanner and listener services maintain logs on S3 for historical tracking and ease of maintenance. These logs are updated after each new message is received or sent.

The data organization on S3 (Figure 2D) is structured based on data types and characteristics. Electrophysiology recordings are grouped by experiment batch, assigned a universally unique identifier (UUID), and paired with a “metadata.json” file for overall content description and experiment notes. We create sub-buckets: “original/data” for raw recordings and “derived/algo” for analysis output, where “algo” represents the algorithm used to analyze the data. Additionally, we maintain a “service” bucket for the chained job scheduler and logging of listener and scanner activities. Since the computing clusters are designed to run containerized data processing jobs, we have created docker images for electrophysiology algorithms with minimum software dependencies.

As illustrated in Figure 2E, when the listener deploys a job to NRP, the platform assigns a node with all the requested resources. The node creates a pod, pulls the docker image from Docker-Hub, and retrieves data from S3 to run the analysis. The processing results are then uploaded back to S3 from the container. Figure 2F demonstrates an example of a containerized batch processing algorithm. In this container, a Python script reads an electrophysiology recording, performs spike sorting on the raw data to identify putative firing neurons (single units), applies autocuration to preserve high-quality units, and generates both visualization figures and spike data for the recording. The spike data is stored as a NumPy data structure with temporal and spatial information of the single units.

Figure 2G shows the Python Kubernetes object configuration in the listener for job execution. This configuration specifies the number of CPUs and GPUs and the amount of memory and storage required to run a specific container. These resource allocations are calculated based on the algorithm workload and data size, optimized for efficient utilization of cloud computing resources. To execute a specific container, the configuration is provided with the corresponding docker image, input data (such as the recording or derived results from the recording), and metadata (including data format or parameter settings). Examples of Kubernetes configurations can be found in Supplementary Table S2.

### This Pipeline Enables Versatile Jobs

Data processing and analysis often require multiple iterations for new experiments due to changes in recording hardware, biological samples, and data requirements. To ensure versatility in data processing jobs, we developed a minimum building block for the pipeline and designed various job execution paradigms.

Figure 3A shows the minimum building blocks of our pipeline. It includes paths to S3 data storage and a containerized algorithm. Each algorithm needs two inputs (data and parameters) and produces one output file with results (processed data, visualization figures, and logs). We store input data and outputs in designated buckets on S3 under each UUID. We keep these parameters in designated sub-buckets (“service/params/*algo*”) on S3, named after each algorithm. Users can pick existing parameters or make new ones on the Dashboard’s “Job Center” page (Figure 3B, Supplementary Figure S3).

**Figure 3:**
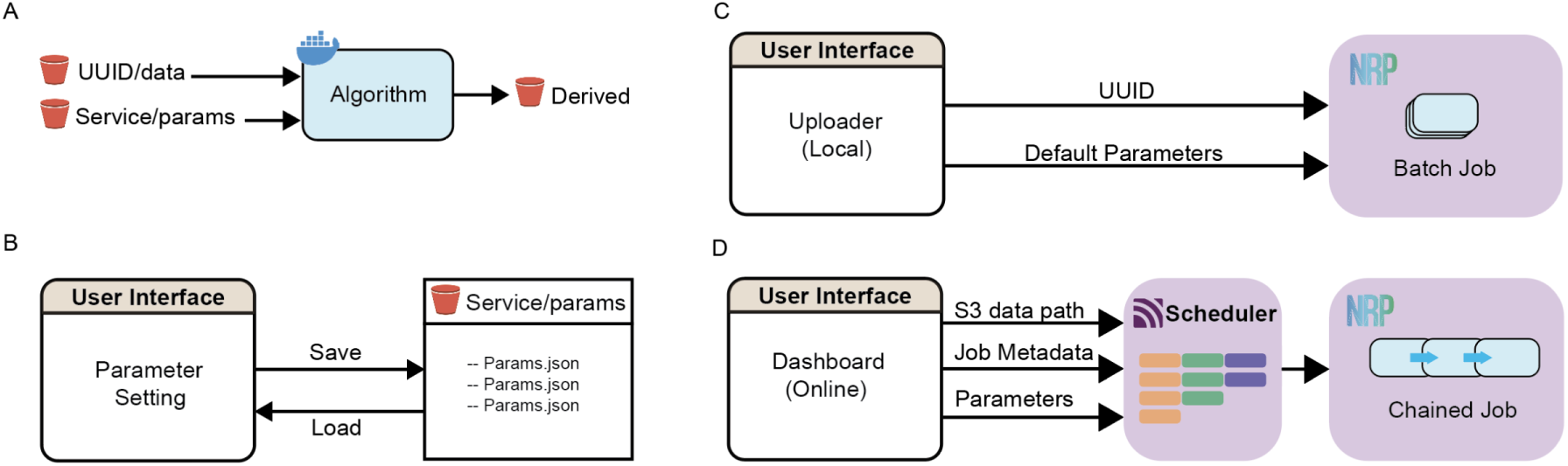
Minimum building block and job types. (A) The minimum pipeline building block utilizing dockerized algorithms and S3 data storage. Data and parameter settings are retrieved from S3, processed by containerized algorithms on NRP, and results are uploaded back to S3. (B) Users can save and load parameter settings to and from the S3 “service” bucket through the Dashboard. (C) Batch processing of numerous recordings is achieved by providing UUID and default parameter settings to the pipeline. Users can initiate this process through the local data uploader. (D) Chained jobs are implemented using a CSV job scheduler containing S3 data paths, job metadata, and parameter settings. Users can initiate job chaining from the online Dashboard.

The pipeline supports both batch processing and chained jobs (Figure 3C,D). Batch processing enables the analysis of numerous recordings using identical parameter settings. All jobs can be processed in parallel on NRP. Users can initiate a batch job from the local data uploader after the experiment. In batch processing, each recording undergoes spike sorting, autocuration, and visualization. Detailed descriptions of these three steps can be found in the Methods section.

As algorithms are packaged in individual docker containers as minimum building blocks, multiple analysis jobs can be chained for a recording, with stage results passed to subsequent jobs upon completion of the previous job (Figure 3D). To implement this functionality, we designed a CSV job scheduler integrated into the Dashboard, Listener, and Scanner services. When users select recordings and a list of analysis jobs from the Dashboard, a CSV file is generated, with each row representing an analysis job. Columns contain sufficient information to initiate the job, including the S3 data path, computing resource requirements (job metadata), and parameter settings. We use the “next_job” column to index the row of the job to run after the current row, allowing for multiple indices. After saving this CSV file to S3, the Dashboard sends a message to the Listener to start the first stage jobs by indexing them in the message body. We create the NRP job name using the CSV file name, enabling the Scanner to differentiate chained jobs from batch jobs by simply parsing the name. Upon completing the first stage jobs, the Scanner sends the Listener an “update” message. The Listener then checks for any available “next_job” in the CSV file and launches the second-stage jobs. Detailed information on job chaining can be found in the Methods section.

### Pipeline Output for Individual Recordings

The pipeline output is designed to be comprehensive, structured, and accessible so the data can be reproduced and distributed easily. Using batch processing algorithms, for example, each processing step produces one compressed file (zip format). For spike sorting, the compressed file is compatible with Phy GUI ^62^. Users can download the file, uncompress it, and open it in Phy to check the sorting result and perform manual curation. We also developed a function to load the data directly into a Python object, enabling automated downstream analysis of the single-unit features. Autocuration, the second step, outputs a compressed file (zip format) containing a spike data object in NumPy array and Python dictionary. This object consists of a spike train list, a neuron data dictionary, the recording’s sample rate, and electrode configuration. The neuron data dictionary has spatial information such as the channel’s coordinates, neighbor channels, and spike features such as waveform and amplitude. The spike train list and the neuron data dictionary index match each other. The size of the autocuration file is approximately 10 times smaller than the spike sorting output by re-constructing the data. For the final step, data visualization, the pipeline generates interactive HTML format figures for the recording and a PNG format figure for each single unit. All of the output files have a log to keep track of the actions and decisions made by the algorithm. To make the data structure consistent, other algorithms’ outputs that are produced by this pipeline are also sorted into NumPy arrays and dictionaries. These outputs can be easily converted to Pandas DataFrame and distributed as tabular data.

Figure 4 illustrates the visualization output for a 10-minute recording from a mouse cortical organoid on day 42 in culture. Figure 4A is a photograph of the mouse cortical organoid on the HD-MEA. Initially, two organoids were plated on the same HD-MEA for this experiment. As the majority of activity originated from the right organoid, our analysis was focused on this organoid. The pipeline’s interactive HTML overview figure includes a footprint map showing the spiking wave-forms on the corresponding electrode locations (Figure 4B). The HD-MEA can detect a unit’s footprint by multiple electrodes and potentially show the neuron’s orientation. Since a single electrode can record activity from many neurons, different colors are used to label the units. Along-side every single unit’s colored footprint (Figure 4B) we provide descriptive electrophysiology features (Figure 4C). We present the unit’s temporal firing rate using 50 ms binning of the spike times over the course of the recording (Figure 4C-i). The result is smoothed by a Gaussian kernel with a sigma of 5. We also provide the amplitude of each spike and a histogram of the amplitude distribution (Figure 4C-ii,iv). Raw spikes and the averaged waveform are also displayed (Figure 4C-iii). Both the amplitudes and raw spikes are from the best channel which recorded the highest mean amplitude of the unit. Interspike interval (ISI) is a crucial feature for neurons, as it is associated with firing patterns and cell types ^23,24,63,64^. We show this information through an auto-correlogram in the range of -50 to 50 ms and a histogram of ISI values in the range of 0 to 50 ms (Figure 4C-v,C-vi).

**Figure 4:**
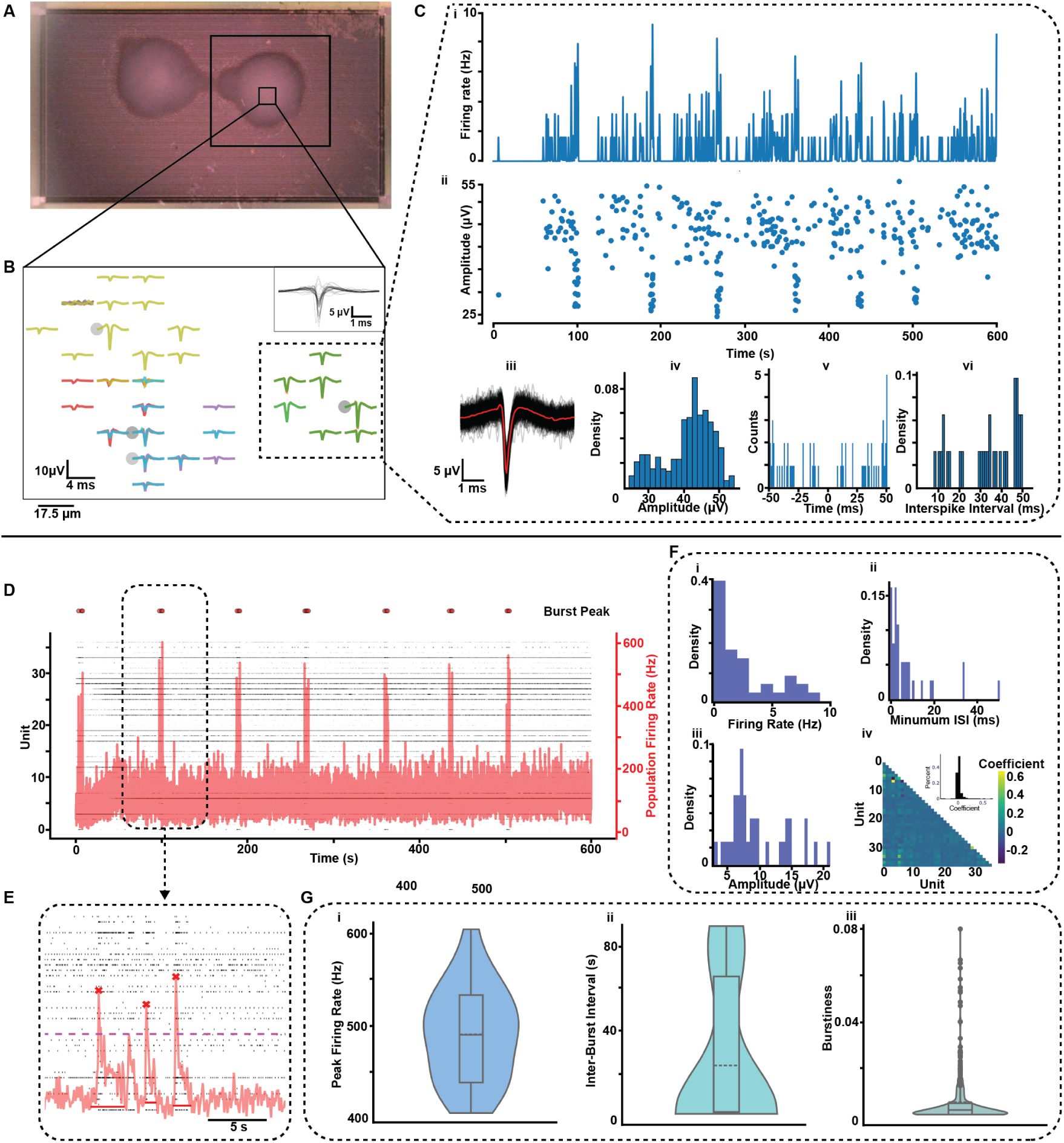
Pipeline Output for an Electrophysiology Recording. (A) Photograph of a mouse organoid on HD-MEA. (B) Zoomed-in view of spiking activities in the mouse organoid. Each color represents a single unit. Waveforms from all single units are shown in the top right corner. (C) Spiking features for the single unit labeled in B: i) Firing rate distribution over the recording time, calculated by binning spike train with a 50 ms time window. ii) Amplitude of each spike over the recording time. iii) Raw spike waveforms (black) and the averaged waveform (red). iv) Amplitude distribution. v) Auto-correlogram from -50 ms to 50 ms. vi) Interspike interval distribution for intervals in the 0 - 50 ms range. (D) Spike raster (black) overlaid with population firing rate (red) for the recording. Dots above the plot label population burst peaks. (E) Zoomed-in view of a population burst. (F) Distribution of i) unit firing rates, ii) minimum ISIs, iii) mean amplitudes, and iv) spike time tiling coefficients. (G) Violin plots showing the distribution of i) population burst peak firing rates, ii) interburst intervals, and iii) burstiness of each single unit.

In addition to the footprint map, the interactive HTML overview figure includes a spike raster and several statistical plots for population features for the organoid. The spike raster shows each unit’s spike times and the population firing rate with labeled burst peaks (Figure 4D,E). Bursts are detected by thresholding the population firing rate. We show burst features such as the distributions of peak firing rate, interburst interval, and each unit’s burstiness index in violin plots (Figure 4G). Furthermore, we display the distribution of firing rates, minimum ISI values, and mean spike amplitudes for all single units in the recording (Figure 4F-i,ii,iii). We also illustrate the pair-wise correlation of units’ firing activity by calculating the Spike Time Tiling Coefficient (STTC) ^65^ value of each unit relative to the others. We designed the overview figures to be interactive, allowing users to zoom in for a closer examination of the data. The figures for individual units are high-resolution. These figures can give users useful information to evaluate the recording object and perform cross-comparisons. Detailed descriptions of data visualization can be found in the Methods section. The complete figures are available in the Supplementary Figure S4, S5.

### Longitudinal Organoid Electrophysiology Properties

Longitudinal neuronal recordings provide invaluable data to study how neuron activity patterns change over time. The cortical organoid shown in Figure 4A was subjected to hourly ten minute recordings on the HD-MEA over seven days (see Voitiuk et al., 2024^51^). During this experiment, recordings were automatically scheduled at the beginning of each hour, uploaded to S3, and processed by the pipeline. Data processing included spike sorting using Kilosort2 and autocuration with quality metrics. Detailed descriptions of the data processing can be found in the Methods section.

Over time, we observed an increasing number of single units and intensified spiking activity. Figure 5A illustrates the time-lapse images of the units’ locations and their action potential amplitudes on the HD-MEA. With a grayscale color bar, the darker color denotes a higher amplitude. The scale ranges from 0 *µ*V (white) to 30 *µ*V (black). There is a noticeable increase in the activity intensity and clustering of active areas as days progress, especially prominent between days 3 to 5. The spike activity across recordings can be found in Supplementary Video SV1.

To visualize the development of the organoid neuronal network we plotted the number of detectable units in each recording (Figure 5C,D) and the individual unit firing rate (Figure 5E,F) over the recording time course. Figure 5C shows the distribution of the unit count across recordings for each day, while figure 5D shows the average number of units each day. There is a substantial growth in the number of units from day 0 to day 2 and decreased variability among the recordings. From day 2 to day 7, the number of units is relatively stable, with an increased variability across the samples. There is a clear upward trend for the firing rate for the individual neural units from each recording (Figure 5E) and the average firing rate for each day (Figure 5F). As the days progress, the firing rate distribution of individual units becomes wider with some units showing higher firing rates while other units have a firing rate between 0 and 10 Hz.

**Figure 5:**
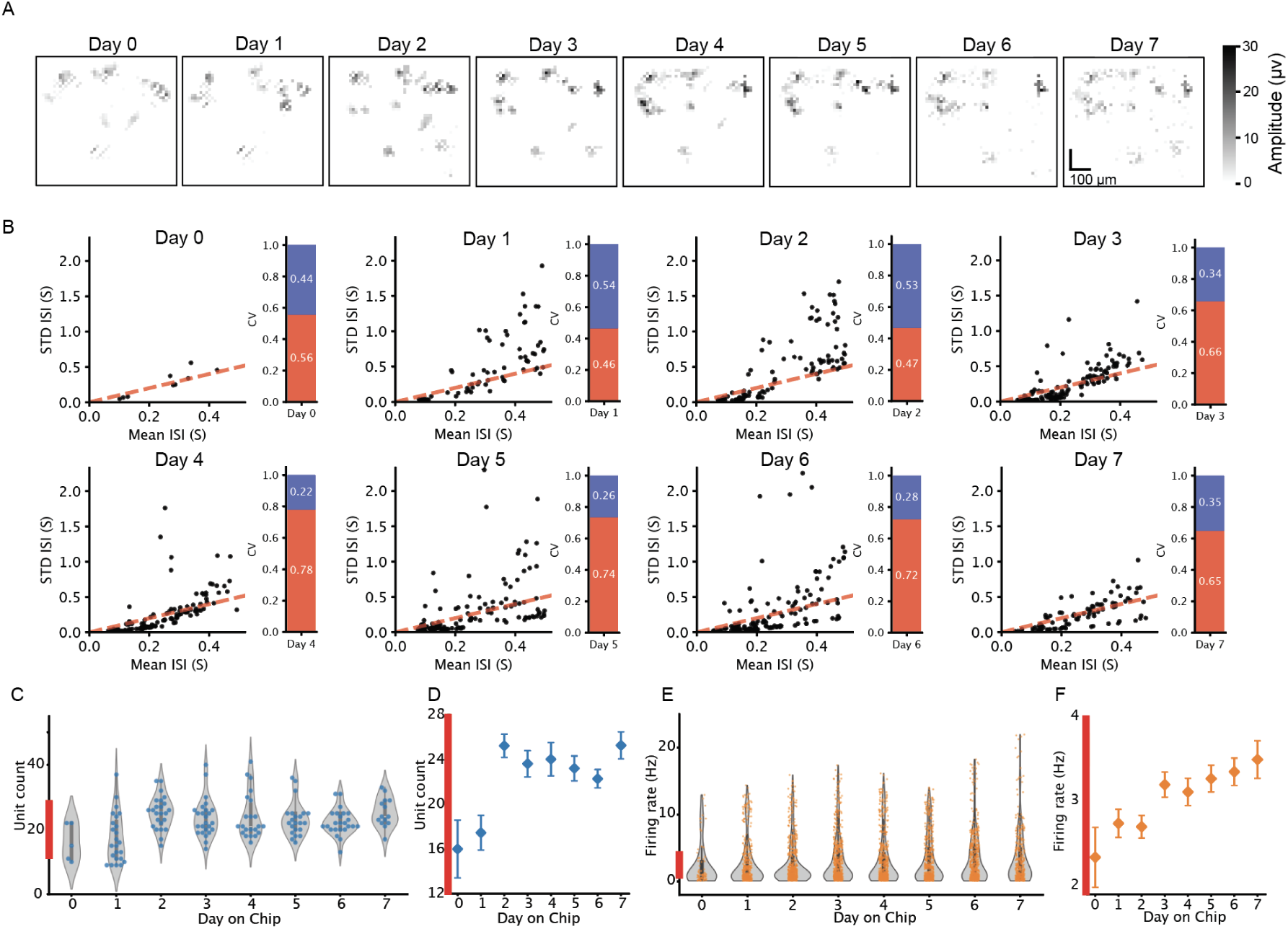
Single neuron features from hourly recordings over days. (A) Spatial area of spiking activity in the mouse organoid on the HD-MEA over the recording time course. Color intensity corresponds to the amplitude of the neuron’s action potential. (B) Changes in the Coefficient of Variation (CV) of interspike interval distribution over time. The bar plot shows the percentage of units with CV < 1 (red) and CV >= 1 (blue). (C) Distribution of the total number of single units for each day. (D) Average unit count with standard error of the mean (SEM) over time (Day 0: 16±2.58, Day 1: 17.45±1.55, Day 2: 25.25±1.03, Day 3: 23.64±1.18, Day 4: 24.04±1.49, Day 5: 23.22±1.12, Day 6: 22.29±0.81, Day 7: 25.28±1.20). (E) Single unit firing rate distribution over the 7 days. (F) Average firing rate (Hz) with SEM over time (Day 0: 2.33±0.35, Day 1: 2.74±0.16, Day 2: 2.7±0.13, Day 3: 3.19±0.14, Day 4: 3.11±0.16, Day 5: 3.27±0.15, Day 6: 3.35±0.16, Day 7: 3.49±0.22)

We also found changes in neuron firing patterns over time. The neural unit firing patterns are represented by the coefficient of variation (CV) of interspike intervals (ISI) ^66,67^. We show the evolution of CV by plotting the standard deviation of ISI to the mean of ISI for each unit over the 7 days. The stacked bar charts represent the proportion of neurons with different CV values, where the red portion indicates neurons with CV < 1, and the blue portion represents neurons with CV*≥*1 (Figure 5B). Over the 7 days, we observe a trend of an increasing number of units showing a more regular pattern with CV < 1, implying the maturity of the neural network. Day 0 to day 2 starts with a fairly even split, with slightly more neurons having CV*≥*1 (44%, 54%, and 53%) and CV < 1 (56%, 46%, and 47%). As time progresses, there is a clear shift towards neurons with CV < 1. By day 6, the majority of neurons have CV < 1 (72%), and a small proportion of neurons have CV*≥*1 (28%).

Overall, this analysis suggests maturation of the mouse cortical organoid neuronal activity over the 7 days with increases in both the number of units detected and their firing rate. The increased firing rate variability could indicate the emergence of more complex and heterogeneous neural circuits within the organoid.

### Neuron Tracking for Longitudinal Recordings

The consistency of the pipeline enables tracking putative neurons throughout the longitudinal experiment, as the same processing steps and parameters are applied to all datasets. A trackable unit can be identified by its consistent spike waveform and location on the HD-MEA. After spike sorting a recording, we gathered the average waveform (2.5ms), the best channel’s location (x, y coordinate on the HD-MEA), footprint, and firing rate for each single unit. We used a waveform clustering algorithm (WaveMap) ^4,24^ to label the units and observed the change of electrophysiological features across multiple days. We ran WaveMap using both the waveform and the best channel’s location. The best channel is defined as the one that recorded the unit’s highest mean amplitude. For each unit, we concatenated the best channel’s location to the end of the wave-form. Then, we aggregated units from all recordings. The waveforms and the locations were normalized separately. As a result, WaveMap yielded 20 distinct clusters for the mouse organoid, as shown in Figure 6A. For each cluster, we characterized the waveform features by measuring the trough-to-peak width and Full Width at Half Maximum (FWHM) of the amplitude. The violin plots (Figure 6C) show significant differences in the waveform features among clusters, indicating potentially different cell types in the organoid. Details of running the algorithm can be found in Methods: Waveform Clustering for Cell Tracking.

**Figure 6:**
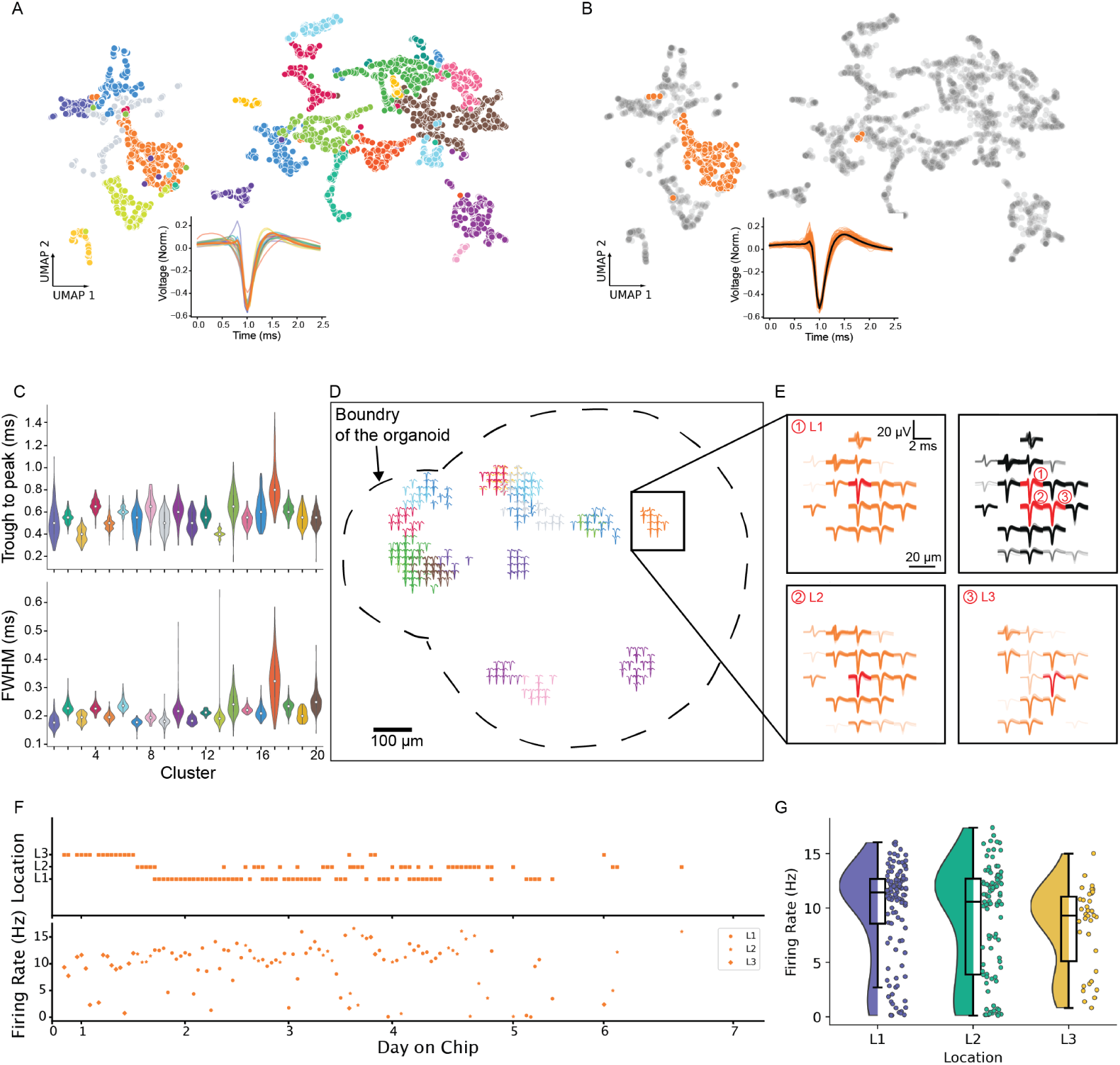
Neuron tracking in a mouse organoid over seven days of recording Pipeline output demonstrating the capability of neuron tracking for longitudinal recordings. (A) UMAP of waveform clusters with location coordinates. The inset shows the mean waveforms of each cluster from a total of 20 clusters superimposed. (B) The cluster of interest (orange) is highlighted on the UMAP, with other clusters in gray. Inset displays the individual waveforms from this cluster, with the mean waveform in black obtained by averaging all the waveforms. (C) Distribution of waveform features for each cluster. The features include trough-to-peak width and Full Width Half Maximum (FWHM) of the amplitude. (D) Footprint projection on the cortical organoid recorded on the MEA. The footprint is color-coded according to the UMAP cluster. Dashed lines show the boundary of the cortical organoid. (E) Footprints for the cluster of interest overlaid across the recording time course. L1, L2 and L3 are the best channels on the footprints. (F) Temporal tracking of location and firing rate change for the units in the orange cluster. (G) Firing rate distribution for the three locations.

For a trackable unit at a static location on the organoid, the unit’s waveforms sampled across recordings should be in the same cluster and appear on adjacent recording channels. Using HD-MEA, we can locate a unit within a small area with an electrode pitch of tens of micrometers (17.5*µ*m for MaxOne HD-MEA). We labeled each footprint by the color of the corresponding waveform cluster (Supplementary Video SV2) and observed the duration and change of its best channel (Supplementary Figure S6) throughout the recordings. For each cluster, we summarized the best channels for each recording and the frequency of each channel (Supplementary Figure S7) that shows activity.

Among these clusters, we selected Cluster 4 as our primary focus (Figure 6B, D). Figure 6B shows this cluster on the UMAP and the waveforms across recordings. We observed the channel locations of the units in this cluster and arranged the footprints from the three adjacent channels that showed the most activity. These activities are highly likely to be from an individual neuron. We labeled the best channels as L1, L2, and L3 and overlaid corresponding footprints for each channel (Figure 6D,E). On an HD-MEA, the electrical signal from a neuron can be picked up by nearby electrodes, which can be beneficial in identifying a neuron’s orientation and movement. During the experiment, this unit initially showed activity on L3. Then its signals were sampled mostly between the two main locations L1 and L2 (Figure 6F). We calculated the firing rate for each sample across recordings and locations (Figure 6F), and grouped the firing rates for each location in Figure 6G. Interestingly, while distributions of firing rates between L1 and L2 did not differ significantly (two-sample Kolmogorov–Smirnov test, p=0.11), there was a significant difference between L2 and L3 distributions (two-sample Kolmogorov–Smirnov test, p=0.019). This finding suggests that L3 may represent a subset of activity of L2 based on differences in their respective footprints.

Using this study, we show the pipeline provides stable, consistent and reproducible data analysis. The neuron tracking function can improve our understanding of an individual neuron’s long-term activity by monitoring its electrophysiological features. Thus, this pipeline offers new possibilities to investigate neural dynamics, plasticity, and neural circuit development.

### Pipeline Applied to Optogenetics Modulation of Epileptiform Activity from Human Hippocampus Slices

Epilepsy is a neurological disorder characterized by abnormal brain activity resulting from an imbalance between excitatory and inhibitory processes ^69^. Light-responsive channelrhodopsins enable optogenetic interventions to modulate the neuronal activity of brain tissues. We applied this pipeline’s data processing and analysis functionality to study the optogenetic modulation of neural circuits from human hippocampus slices.

Before optogenetics illumination, human organotypic tissue slices from hippocampus tissue were established. The hippocampus tissue was obtained from patients with drug-refractory temporal lobe epilepsy, sliced to 300*µ*m, and cultured at an air-liquid interface. Slices were transduced with AAV9 carrying an HcKCR1 transgene driven by a CaMKII*α* promoter and a fluorescent tag (eYFP) (see Andrews et al., 2024^68^). A hippocampus slice was plated on an HD-MEA (Max-One) for electrophysiology recording, with a fiber-coupled LED to illuminate the slice from the top. Since HcKCR1 encodes a kalium channelrhodopsin, a light-gated potassium channel that hyperpolarizes the neuronal membrane, the probability of neuronal spiking is reduced when activated by 530nm light. Bicuculline was applied to the slice after plating to increase neuron firing rates, inducing epileptiform activity. During the experiment, we illuminated the slice for 10 seconds at 0.6 light intensity (35.8 mW/mm2) of the LED driver, and observed the neuronal population firing activity for 10s prior to illumination (Pre-Stim), 10s during illumination (Light-On) and 10s following the end of illumination (Light-OFF). The experimental setup is shown in Figure 7A. Each HD-MEA recording was processed by the described pipeline using spike sorting and autocuration algorithms. Neuronal activities were aligned to optogenetics stimulation timestamps that were synchronized with the recording. As illustrated in Figure 7B, the units’ footprints were overlaid with NeuN staining of the slice on the HD-MEA recording area, showing the physical location of the spiking activity. Examples of spike waveforms and auto-correlograms are shown.

**Figure 7:**
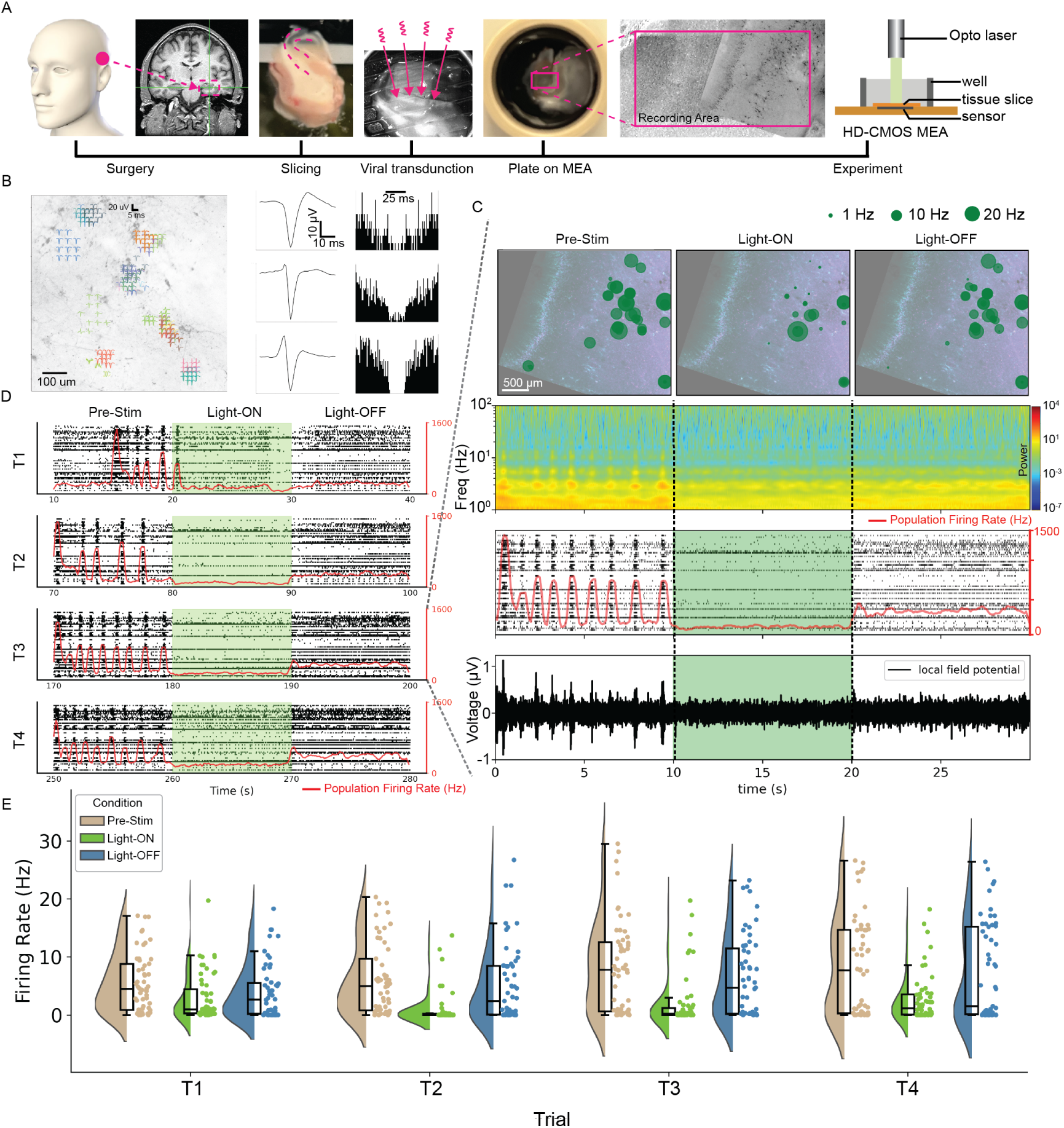
Pipeline facilitates seizure study by analyzing electrophysiology data from human hippocampus brain slice with optogenetic stimulation. (A) Experimental setup for hippocampus slice HD-MEA recording ^68^. Brain tissue from a seizure patient in 300*µ*m thick slices cultured expressed channelrhodopsin delivered through adeno-associated virus (AAV) delivery. The slice is placed on HD-MEA for simultaneous optogenetic stimulation and recording detailed in Andrews et al., 2024^68^ (B) NeuN-stained hippocampus slice overlaid with an image of the slice on HD-MEA and footprint of spiking activities on the slice. Example spike waveforms and auto-correlograms from representative neurons. (C) Hippocampus slice overlaid with single units’ firing on the HD-MEA. The change in firing activity is shown for the three steps of Trial 3 (T3). From top to bottom, the panels display the single unit’s location overlaid with GFP-stained slice, firing rate, local field potential spectrum, spike raster, and voltage data from all recording channels. (D) Spike raster with population firing rate for four experimental trials under Pre-Stim, Light-On, and Light-Off conditions. The population firing rate shows epileptiform activity suppressed by optogenetics illumination, with the firing rate remaining low afterwards. (E) The single unit’s firing rate distribution for each Trial in (D).

The optogenetics modulation of population firing is shown in Figure 7D and Supplementary Figure S8. The bicuculline-provoked recurrent burst activity was rapidly suppressed by the illumination. Interestingly, the burst activity didn’t completely recover when the illumination was off. The firing rate suppression of individual neuronal units was consistent among trials (Figure 7E). This pipeline is capable of providing multiple perspectives of an individual neuron’s firing activity (Figure 7C). In addition to firing rate, the suppression of activity is visualized on the hippocampus slice through the electrodes on the HD-MEA. The pipeline can extract local field potential (LFP) data by applying a 5th-order Butterworth bandpass (0.1-100Hz) filter to the raw voltage data. During the 10 seconds “Light-ON” period, activities in the LFP frequency bands were also attenuated.

This application showcases the pipeline’s adaptability to diverse experimental paradigms, extending its utility beyond basic neural activity analysis to more complex neurological disease studies. This advancement opens new avenues for studying neurological disease mechanisms and potentially developing therapeutic approaches, highlighting the pipeline’s significance in translational neuroscience research.

## DISCUSSION

The cloud-based electrophysiology data pipeline in this study represents an advancement in the processing, analysis and visualization of HD-MEA recordings, which are enabled by IoT and cloud computing technology. The integration of the MQTT messaging protocol enables automation and communications between various components of the pipeline as well as user interfaces. The cloud-based infrastructure provides storing and parallelized processing capabilities for large volumes of longitudinal, high-throughput experiments. Furthermore, the flexible and modular architecture can meet different data analysis goals, providing comprehensive electrophysiology feature extraction and making this pipeline scalable.

### Limitations and Adaptability

The reliance on IoT messaging and cloud computing resources may present challenges to laboratories with limited access to these services or those working with sensitive data that requires local processing. While containerization provides excellent modularity, it may introduce overhead in terms of deployment complexity.

The pipeline’s adaptability, however, offers significant opportunities for scalability and customization. By applying the IoT protocol, this pipeline can access data from multiple devices across different locations and communicate with users simultaneously. Since both the algorithms and Dashboard are containerized, they can be configured to run on either High Performance Computing clusters or local hardware. Processing speed may be limited because of lacking parallelization and computing resources. The pipeline is adaptable to various recording platforms and neural interfaces that incorporate the NWB data format, such as Neuropixels ^70,71^. Furthermore, the open-source code allows laboratories to customize the pipeline to meet their data requirements.

### Consistent and Reliable Data Processing

By containerizing and using the same parameter settings for algorithms across all recordings, we ensure that data is processed uniformly without human intervention. This consistency is essential for longitudinal studies, where analyzing changes in neural activity over time requires a stable and reproducible processing framework ^30,72^. Containerization enables the algorithms to run independently of the operating system, reducing the effort to re-build and test algorithms across different software environments. The autocuration algorithm significantly reduces the manual labeling effort for recordings containing hundreds of neurons, while still providing high-quality data. It eliminates labeling variations caused by human bias from differences in experience and research background. In addition, we minimized the effort required to maintain parameter settings by storing configurations on S3 and accessing them through the user interface. Batch processing also simplifies algorithm reconfiguration to meet the change of data requirements.

### Data Organization and Visualization

To ensure straightforward and efficient data organization, we implemented a hierarchical structure with a strategic naming convention for Ceph/S3 buckets. This is complemented by metadata files that store detailed experiment-related information and log files for data processing. The integration of user interfaces, such as the data uploader and Dashboard, provides researchers with tools to access and visualize their data. The Dashboard, built using the Plotly Dash library, includes interactive features that allows researchers to explore and analyze their data in depth. This approach has been successfully implemented in recent user interfaces ^73–75^ for visualizing electrophysiology data across multiple scales. By combining data storage, processing and visualization into one multiscale pipeline, we provide researchers deeper insights into complex datasets by decomposing raw data to actionable analyses.

## CODE AVAILABILITY

This electrophysiology data pipeline is an open-source project. The code will be released on GitHub upon manuscript publication and is currently available upon request.

## DATA AVAILABILITY

No new data was generated for this paper. All datasets described were obtained from Voitiuk et al., 2024^51^ and Andrews et al., 2024^68^ (DANDI:001132/0.241119.1801).

## METHODS AND MATERIALS

### Mouse Cortical Organoids

The data presented in this manuscript was collected using an integrated system for neuronal cell culture ^51^. Mouse cortical organoids were made using a protocol described in Park et al., 2024^76^. Cortical organoid recordings were performed on a MaxWell Biosystems MaxOne CMOS HDMEA chip. The system captured 10-minute recordings every hour for seven days. The recording configuration remained consistent throughout the experiment. Details can be found in Voitiuk et al., 2024^51^.

### User Interfaces, Data uploader, and Dashboard

To give researchers full control over their experimental data, we designed user interfaces that enable data management, data processing, algorithm parameter configuration, and result visualization. We developed the data uploader installed on a local computer of the recording device as well as an online Dashboard for remote data access.

The data uploader, created using the Python PyQt library, facilitates the uploading of experiment recordings from a local laptop to S3. Upon opening this application, users can navigate to a folder where recordings are stored. An initial Universally Unique Identifier (UUID) is generated using the date of the recordings, and users can add more descriptive labels to this UUID. Before uploading, users must generate a metadata file by loading a template and inputting any experiment-related information such as notes, cell lines, media used for culture, and recording features for each dataset. Recording features such as recording length, number of active channels, and data path are automatically populated in the metadata template. When users press the upload button, all recordings in the selected local folder are reorganized according to the S3 file structure and are uploaded to the UUID folder on S3. Users can start the data analysis pipeline after uploading by sending a request a message to the job listener.

The Dashboard was created using the Python Plotly Dash library. This library uses callback functions to achieve user-interactive features like dropdown lists, buttons, and tables. We built a multipage website, with each page serving a different purpose.

On the “Data Processing Center” page, users can choose recordings from the S3 dropdown list, select preferred data processing jobs, set parameters, and start NRP jobs. It allows users to perform batch processing or chained tasks for chosen recordings. Batch processing is the most commonly used case since all parameters and algorithms are usually the same for an experiment. Chained tasks are practical for testing parameter and algorithm combinations for new experiment setups. Supplementary Figure S3 shows the job center webpage.

On the “Status” page, users can monitor the job status of current tasks on the NRP cluster. By using the “Show Status” button, users can check jobs labeled with prefix “edp-”. This function parses the information returned by Kubernetes Python API for all jobs in the namespace. Upon refreshing with the button, the NRP job name, running status, and data summary will be displayed on the webpage.

The “Visualization” page is designed to display figures of post-processed results. Users can select a processed recording from the dropdown menu to display an interactive raster plot and electrode map. Clicking a unit on the electrode map highlights its spike times in light red on the raster plot and shows its waveform and interspike interval histogram. This page allows users to evaluate MEA recordings effectively.

### Cloud Data Storage and Organization

Efficient cloud data organization is crucial for optimizing access performance and storage management. In this pipeline, we employ a hierarchical structure with strategically named buckets. We use a UUID that reflects the experiment date and key information. Upon data uploading, the metadata file is automatically generated and uploaded with the raw data. For each UUID, we keep the raw data in “/original/data”, and the processed result files in “/derived/*algo*”, where “*algo*” can be “kilosort2”, “connectivity” and others that are named after the specific algorithms.

We store logging files on S3 for MQTT services and data analysis jobs. Detailed logs provide a comprehensive record of each pipeline component by capturing essential information. These logs enable researchers to track the progression of data processing, identify potential bottle-necks, and troubleshoot issues effectively. Service logs are generated when the MQTT broker sends or receives a message and updated to the S3 “/service” bucket on a schedule. Logs from the data analysis jobs include processing steps, quantities, and malfunctions. These log files are kept in the algorithm output directory.

### Cloud Orchestration

Each analysis algorithm is packaged into a Docker container with the minimum required dependencies, enabling parallel processing of a large volume of electrophysiology recordings in a cloud-agnostic environment. This approach simplifies the addition of new analysis algorithms to the pipeline.

We use Kubernetes to deploy and monitor data processing jobs on NRP. For each container, based on the input data size and algorithm requirements, we request computing resources from NRP, such as the number of CPUs, GPUs, memory, and storage. Supplementary Table S2 summarizes the computing resource requirements for each algorithm on a 10-minute HD-MEA recording with 1000 active channels. When a job is deployed to NRP, a pod with a job is created to run the data in a container. To get the status of the pod, we extract metadata from the return of the Kubernetes “list_namespace_pod” function. From the metadata, we provide the status of the job, such as “running” or “succeeded,” and the timestamps for running this job.

### MQTT Messaging Application

To enable remote job execution for a large number of recordings, we implemented services using MQTT messaging. This infrastructure has been previously described ^47,51^. All messaging services are hosted on the Braingeneers UCSC Genomics Institute server. We package these services into Docker containers and manage them using Docker Compose.

#### Job Listener

We designed a centralized MQTT service to parse analysis job run messages. This service subscribes to specific topics and responds by running the corresponding Docker container on the given data. We assign the topics “experiments/upload” for batch processing or “services/csv_job” for chained tasks.

The message body is designed according to the different topics. For “experiments/upload”, we use the UUID and recording file name from the metadata.json file. The service can assemble the S3 file path for each recording from this information. The computing requirements for running batch processing jobs are written to a look-up table in the listener service.

For “services/csv_job”, we first create a CSV file where each row contains the UUID, recording file name, job type, and computing requirements for running the analysis. We name the CSV file using the current timestamp, upload it to the S3 services bucket, and create a message containing the path to the CSV file and the indices of the CSV rows. When the listener receives this message, it pulls the information from the CSV file and deploys jobs using the given indices. Examples of the messages are included in Supplementary Materials.

#### Job Scanner

When running analysis jobs on the NRP, we use the prefix label “edp-” in the job name. We name batch processing and chained jobs differently. For batch processing, we name the job using a prefix and the recording file name. For chained jobs, we name the job using a prefix, the CSV file name, and the index of the CSV row. This naming convention allows us to parse the job name to determine which analysis algorithm is running on which data.

The job scanner is designed to scan the “edp-” jobs on a schedule with two main aims. First, it notifies the job listener when the current step in a chained job is finished. This message body contains the keyword “update”. When the listener receives this message, it checks the corresponding CSV file to launch any pending jobs related to the current job. The scanner scans NRP every 2 minutes to minimize delays in running chained jobs. Second, it notifies users of their job status via a Slack channel using the messaging bridge service 18. These notifications are sent every 30 minutes.

Job information is pulled from NRP using Python-Kubernetes functions. We use the “list_namespaced_pod” function to get all “edp-” jobs. We loop through them, extracting job name, data file name, job type, and timestamps. This information is stored in a Python dictionary and updated when the scanner service scans NRP on schedule.

The scanner identifies job status and sends messages to other MQTT services. For batch processing jobs, the scanner sends a user notification message when the job status is “pending”, “running”, “failed”, or “succeeded”. Since “failed” and “succeeded” jobs are finished, the scanner removes these jobs from NRP and the dictionary after sending the message. For chained jobs, when a job is finished as “succeeded”, in addition to sending a user notification, the scanner also sends an “update” message to the listener to run the next job.

#### User Notification to a Slack Channel

Both the job listener and scanner can send user notifications. When a run job message is sent to the listener, and the listener successfully deploys jobs to NRP, a notification is sent to the Slack channel with the S3 data path, job type, and “start” status.

To make user notifications human-readable and clear, when the scanner sends messages to the Slack channel, it groups the jobs by UUID and lists the recordings in each UUID.

### Spike Sorting

Spike sorting is fundamental in analyzing extracellular recordings for assigning action potentials picked up by electrode channels to neurons in an ensemble. For the HD-MEA recordings, Kilosort2 was used to sort the raw voltage data into single unit activity. Since HD-MEAs can record one neuron from tens of channels, it is common for spikes from many neurons to overlap in time on a single channel. The template matching and clustering algorithm in Kilosort2 can distinguish spikes between different neurons based on their waveform. Before spike sorting, the raw data is bandpass filtered using an IIR filter with a 300 - 6000 Hz bandwidth. The data type is converted to int16 for running Kilosort2. The voltage detection threshold of Kilosort2 is set to 6 RMS over the baseline. Parameter settings for Kilosort2 are shown in Supplementary Table S1. Spike sorting was performed on the National Research Platform computing cluster with an NVIDIA A10 GPU. The sorting output is saved to a compressed file (zip format) and uploaded to S3. The output file structure is compatible with the software Phy for performing manual curation. An autocuration process is built on top of the sorting result.

### Autocuration

The autocuration process is applied after spike sorting for each single unit. To assess data quality and retain good units for downstream analysis, we evaluate each unit by calculating the Interspike Interval (ISI) violation ratio, Signal-to-Noise ratio (SNR), firing rate, and spike waveform.

We use the curation module from SpikeInterface API ^30^ in our Python script. For ISI violation, we apply the Hill method ^77^ of false positive errors with an absolute refractory period of 1.5 ms. We set the maximum violation rate to 20%. The SNR is calculated using the spike amplitude of a unit and the baseline voltage, with a minimum SNR threshold of 5 RMS. The unit’s firing rate is defined as the total number of firing events divided by the recording length in seconds. In our default autocuration algorithm, this threshold is set to 0.1 Hz.

To check the spiking waveform for a unit, we run the WaveformExtractor class across all active channels and take the average of a maximum of 500 spikes. We then find the best channel and a maximum of eight neighboring locations on a 3x3 grid by sorting their waveform amplitudes on each channel from highest to lowest. The best channel is defined as the channel that captures the neuron’s highest mean amplitude among all recording channels. Since HD-MEAs can record one neuron across multiple channels simultaneously, we expect the waveform distribution to have an adequate layout. This layout is defined as the unit’s footprint. Thus, we save units that show a waveform on the best channel and at least one neighboring channel within a distance of 17.5*µ*m for further analysis.

### Visualization of Electrophysiology Features

For each recording, the pipeline generates an interactive overview figure in HTML format that includes the activity map of the MEA, the neuron’s footprint at its physical location on the map, a spike raster with population burst detection ^7^, and a summary of electrophysiology features for all single units. The population firing rate is smoothed using moving average (20 ms window size) applied to aggregated spike trains, then further smoothed using a Gaussian kernel with sigma = 20. The population burst detection threshold is set to 2 RMS of the population’s baseline firing rate. Burst detection is performed using *scipy.signal.find_peaks* with a minimum peak distance of 800 ms. Burst edges are defined as points where the firing rate drops by 90% from the peak on both sides. The “burstiness index” ^78^ of a single unit is represented by a number from 0 to 1 that measures the synchronization in spiking activity by binning (40 ms bin size) the spike train. Electrophysiology features include interspike interval (ISI), minimum ISI, firing rate, amplitude, spike time tiling coefficient (STTC) ^65^ and average spike waveforms. Distributions of these features for all single units are provided in the interactive figure. Each single unit is also paired with a PNG format figure showing the unit’s footprint, raw and average spike waveform, auto-correlogram (ACG), ISI distribution, instantaneous firing rate and amplitudes. Both the interactive figure and single unit figures are created using the Plotly Python graphing library. Parameters for the visualization are adjustable in the source code.

### Waveform Clustering for Cell Tracking

Neuronal cell types and their spiking waveforms are known to be correlated. To demonstrate the capability of tracking units in longitudinal recordings, we performed waveform clustering using the WaveMAP Python package ^4,24^. This package combines non-linear dimensionality reduction (UMAP) with the Louvain clustering method.

To prepare the waveforms, we first extracted the spike times for each single unit through spike sorting. Since the spike time represents the peak of each spike, we initially took a 5 ms window of the complete waveform from the best channel, then averaged across up to 500 spikes. Before input into the WaveMAP algorithm, we centered the waveforms at their peak and truncated them to 1 ms before and 1.5 ms after the peak. Units with positive spikes were not included in this clustering due to the high possibility of axonal signals. We extracted waveforms for each recording, stacked them into a NumPy array, and pre-processed them using l2 normalization. The total number of waveforms was 3526 from 160 recordings.

Given that the mouse organoid recordings were taken hourly across seven days, and neurons can migrate during development, we appended the corresponding electrode location (x, y) to the end of each waveform for clustering. The location was normalized as a percentage of the maximum x and y coordinates, respectively, to ensure the data range was within [0, 1], comparable to the normalized waveform.

The UMAP parameters were set with 20 neighbors and a minimum distance of 0.1, while the Louvain clustering resolution was set to 1.5. As a result, the algorithm identified 20 distinct clusters and assigned a color to each. Based on the clustering results, we plotted the footprint of each unit on the electrode map using the assigned color. Throughout the recordings, we were able to track changes in a neuron’s location and firing rate.

### Local Field Potential

Local field potentials (LFPs) are low frequency signals up to about 500 Hz that are generated by multiple signal processes in a neural population ^79^. These signals are traditionally decomposed into frequency domain. In this pipeline, we focused on LFPs in the range of 0.1 to 100 Hz, and subband frequencies as delta (0.5 - 4 Hz), theta (4 - 8 Hz), alpha (8 - 13 Hz), beta (13 - 30 Hz) and gamma (30 - 50 Hz).

To get LFPs and subband frequencies, we first bandpass filter the raw voltage signal from all recording channels with 0.1-100 Hz 5-order Butterworth filter. Then, these signals are downsampled 1 kHz. A second bandpass filter is applied to separate subband frequencies. We use spectrograms to show the signal strength of different subbands. A spectrogram is the time-frequency spectrum of the local field potential signal, based on the power values, over the given time and frequency range. We applied a continuous wavelet transform (CWT) on the local field potentials to obtain wavelet coefficients and corresponding frequencies using the complex Morlet wavelet (’cmor1-1’) in PyCWT library. Signal strength is computed as the magnitude squared of the wavelet coefficients and smoothed using a Gaussian filter with sigma of 2.

## ACKNOWLEDGMENTS

This work was supported by the Schmidt Futures Foundation SF 857 and the National Human Genome Research Institute under Award number 1RM1HG011543 (D.H., S.R.S and M.T.), the National Institute of Mental Health of the National Institutes of Health under Award Number R01MH120295 (S.R.S.) and 1U24MH132628 (M.A.M.-R. and D.H.), the National Institute of General Medical Sciences (NIGMS) under award number K12GM139185 and the Institute for the Biology of Stem Cells (IBSC) at UC Santa Cruz, the National Science Foundation under award number NSF 2034037 (S.R.S and M.T), and NSF 2134955 (M.T., S.R.S and D.H). This work was supported in part by National Science Foundation (NSF) awards CNS-1730158, ACI-1540112, ACI-1541349, OAC-1826967, OAC-2112167, CNS-2100237, CNS-2120019, the University of California Office of the President, and the University of California San Diego’s California Institute for Telecommunications and Information Technology/Qualcomm Institute. Thanks to CENIC for the 100Gbps networks. H.E.S. is partially supported by the Graduate Research Fellowship Program of the National Science Foundation. The authors want to give special thanks to Anna Toledo, Lon Blauvelt, Catharina Lindley, the IBSC Cell Culture Facility (RRID:SCR 021353), National Research Platform (NRP), and the UCSC Life Sciences Microscopy Center (RRID:SCR 021135) for valuable resources and assistance.

## AUTHOR CONTRIBUTIONS

J.G., K.V., D.F.P., and A.R. conceived the project and established the cloud storage organization, cloud computing, and IoT codebase. J.G. designed the pipeline architecture, built the pipeline components, contributed to data collection, analyzed and interpreted the data, created figures and wrote the manuscript. A.R. developed the data uploader and contributed to MEA recordings and spike sorting. A.S. assisted with data analysis code. J.L.S. provided insights into pipeline design, interpreted the data, and contributed to data visualizations. R.C. contributed to the conceptualization of the cloud-based data architecture. S.H. and H.E.S. cultured and provided mouse cortical organoids for MEA recording. K.V. and S.T.S. designed and conducted mouse cortical organoid experiments, provided data, and interpreted the data. E.F.C. provided human tissue samples. J.P.A. obtained human tissue samples, designed and conducted experiments, provided data, interpreted the data, and performed histology. J.P.A., J.G., K.V., A.R., and M.A.T.E. conducted optogenetics experiments, gathered data, and interpreted the results. T.J.N., M.A.M.-R., D.H., T.S., S.R.S., and M.T. provided mentorship, intellectual consultation, input on experimental design and analytic methods, discussed the results. M.A.M.-R., D.H., S.R.S., and M.T. provided funding for the project. All authors commented, edited, and approved the manuscript.

## DECLARATION OF INTERESTS

K.V. and S.T.S. are co-founders and D.H., S.R.S, M.T. are advisory board members of Open Culture Science, Inc., a company that may be affected by the research reported in the enclosed paper. All other authors declare no competing interests.

## Notes

### Competing Interest Statement

K.V. and S.T.S. are a co-founders and D.H., S.R.S, M.T. are advisory board members of Open Culture Science, Inc., a company that may be affected by the research reported in the enclosed paper. All other authors declare no competing interests.

### Summary of Updates

Figure 6 updated color. Added citations to the Introduction.

