## Supplementary Material for "Multiscale Cloud-Based Pipeline for Neuronal Electrophysiology Analysis and Visualization"

#### Flowchart for the Listener

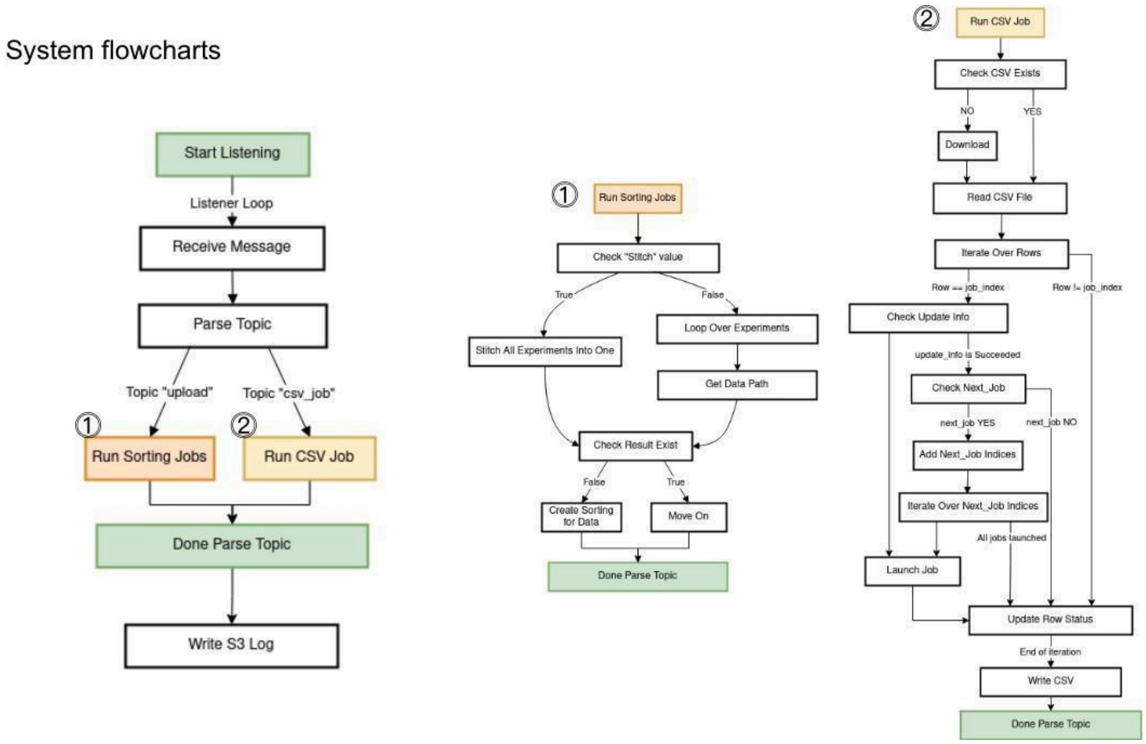


**Figure S1: Breakdown of the MQTT job listener.**

**Table S1. Kilosort2 parameter setting**

| | | | |
|---|---|---|---|
| detect_threshold | projection_threshold | precluster_threshold | car |
| 6 | [10, 4] | 8 | 1 |
| minFR | minfr_goodchannels | freq_min | sigmaMask |
| 0.1 | 0.1 | 150 | 30 |
| nPCs | ntbuff | nfilt_factor | NT |
| 3 | 64 | 4 | 65600 |
| keep_good_only | total_memory | n_jobs_bin | trange |
| False | 2G | 64 | [float(0), float('inf')] |

**Table S2. Kubernetes Job Computing Resource Requirements**

| Analysis Job | CPU Request | GPU Request | Memory Request | Disk Request |
|---|---|---|---|---|
| Batch Processing | 12 | 1 | 32 | 400 |
| Spike Sorting | 12 | 1 | 32 | 400 |
| Autocuration | 8 | 0 | 32 | 400 |
| Visualization | 2 | 0 | 16 | 8 |
| Local Field Potential | 4 | 0 | 64 | 64 |

#### Examples of MQTT messages

An example MQTT message in JSON format to run batch processing jobs.

{'uuid': 'yyyy-mm-dd-e-pipeline-test',
'overwrite': True,
'ephys_experiments': {'yyyy-mm-dd-Thhmmss-chip00000-1':
{'blocks': [{'path': 'original/data/'yyyy-mm-dd-Thhmmss-chip00000-1.raw.h5'}]},
'yyyy-mm-dd-Thhmmss-chip00000-2’:
{'blocks': [{'path': 'original/data/'yyyy-mm-dd-Thhmmss-chip00000-2.raw.h5'}]},
}}

An example to run individual analysis jobs through a CSV scheduler.

{'csv': 's3://braingeneers/services/mqtt_job_listener/csvs/yyyymmddhhmmss.csv',
'update':
{'Start': [$N_1$, $N_2$, $N_3$, $N_m$, $N_n$]},
'refresh': False}

Table S3 shows the content in the CSV file for the message to run correspond jobs.

**Table S3. CSV Job Scheduler**

| | |
|---|---|
| index | N |
| status | ready |
| uuid | yyyy-mm-dd-e-pipeline-test |
| experiment | yyyy-mm-dd-Thhmmss-chip00000.raw.h5 |
| image | creator/docker_algo: v0.1 |
| args | python run.py |
| params | algo_default_params.json |
| cpu_request | 12 |
| memory_request | 32 |
| disk_request | 400 |
| GPU | 1 |
| next_job | None |

#### Figures for the User Interfaces

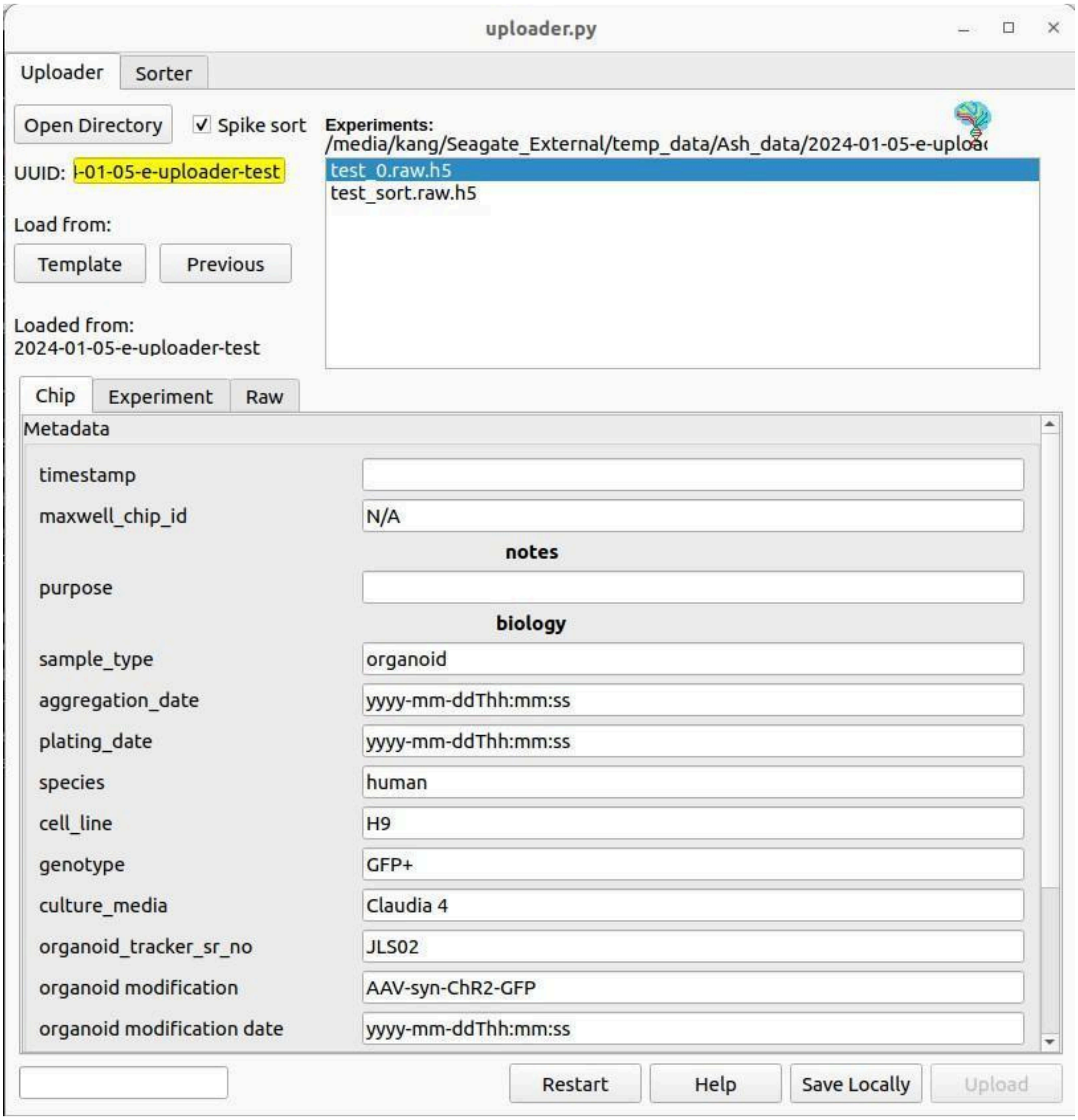


**Figure S2: Maxwell Data Uploader**

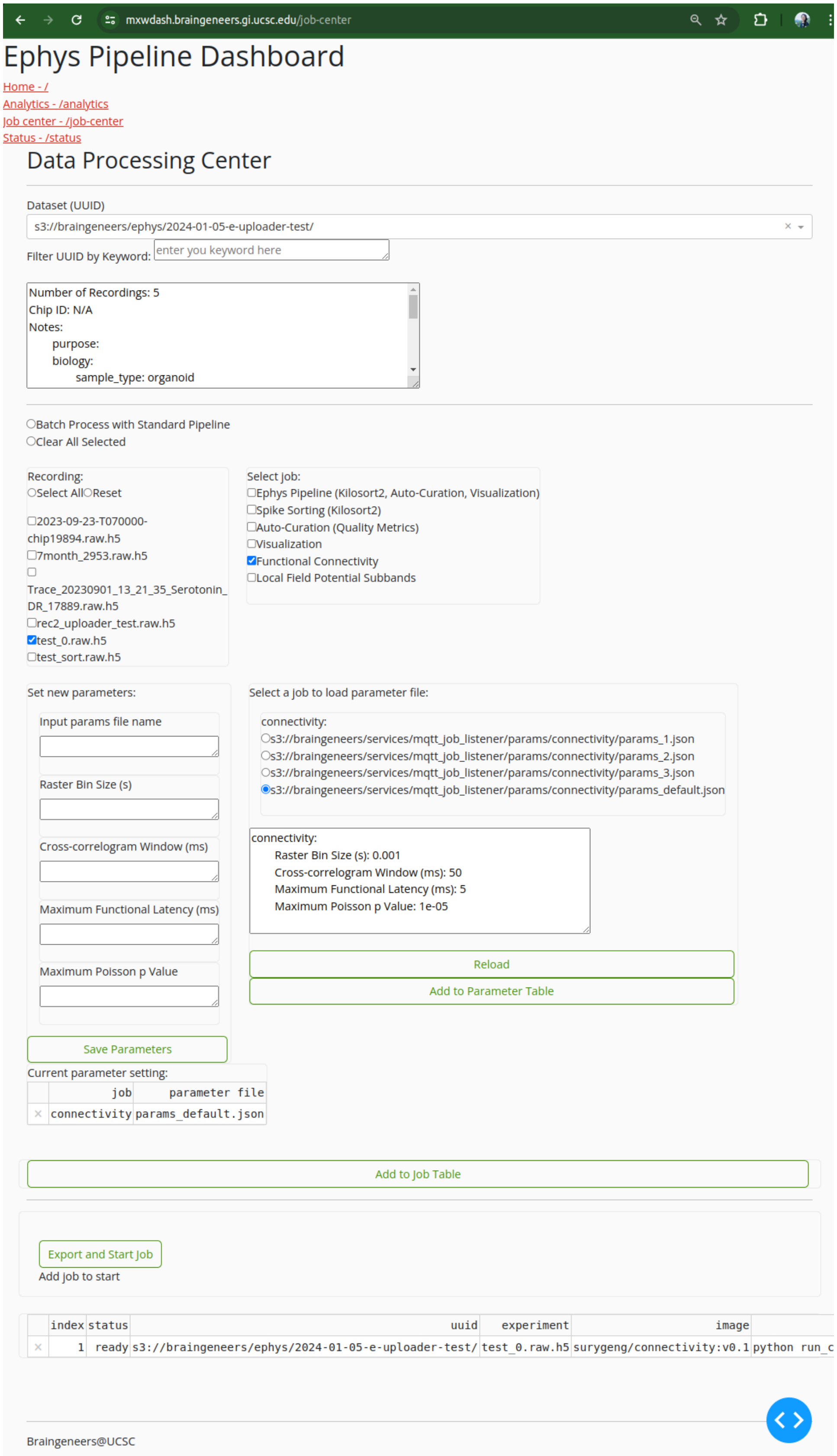


**Figure S3:** Maxwell Dashboard, Job Center

### Pipeline Output: Visualization

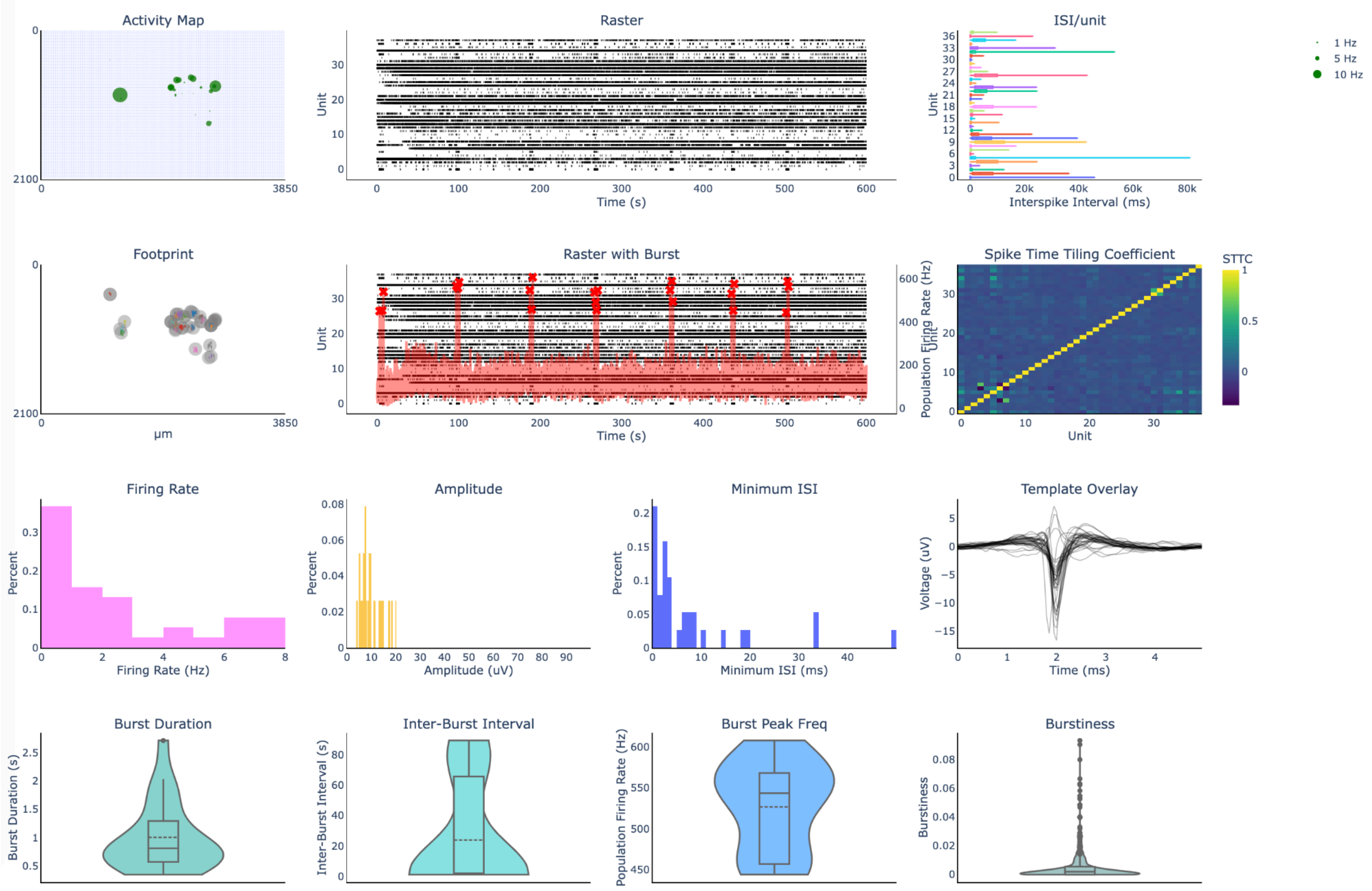


**Figure S4:** **Overview Figure of a recording.**

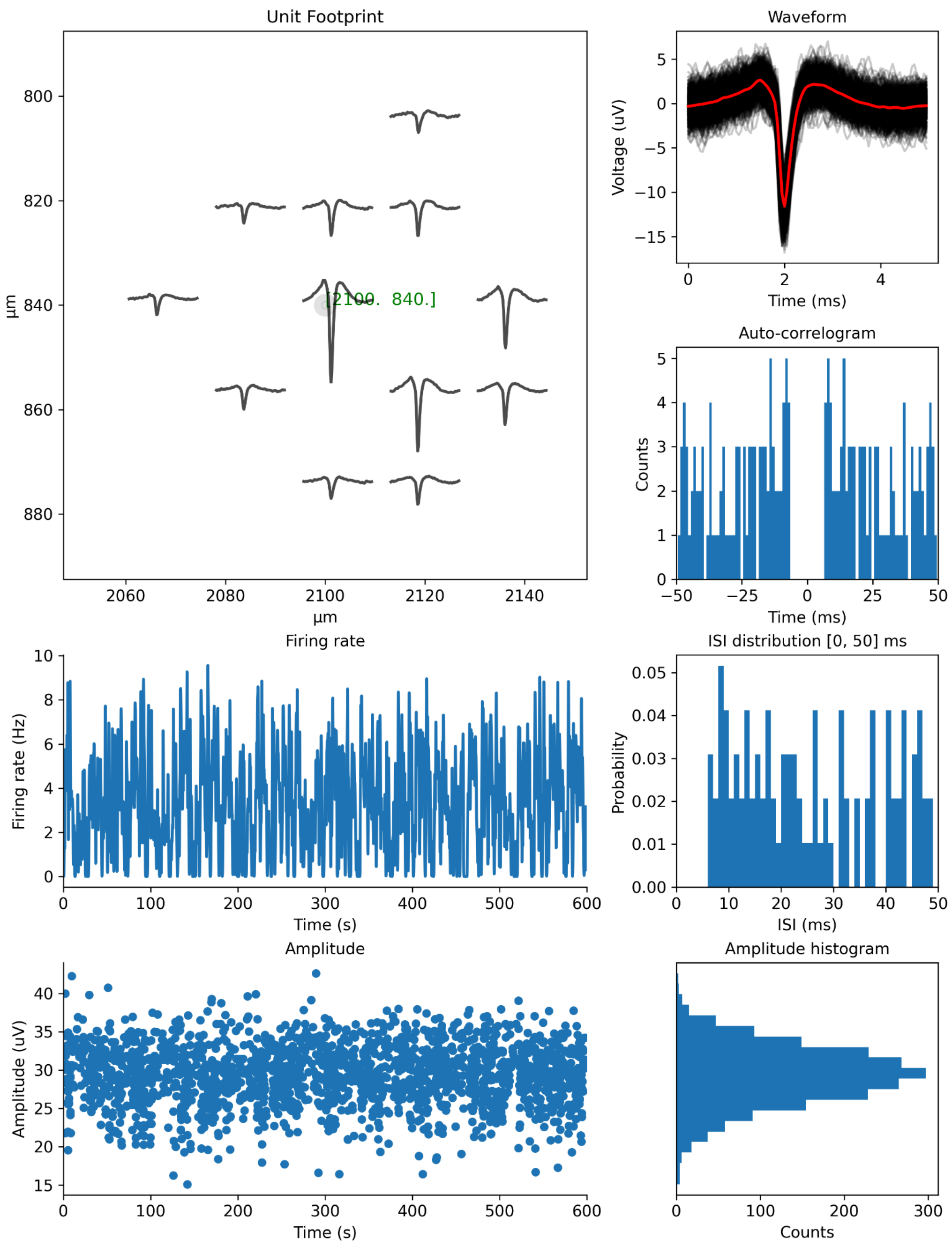


**Figure S5: Electrophysiology features of a single unit.**

**Waveform Clustering**

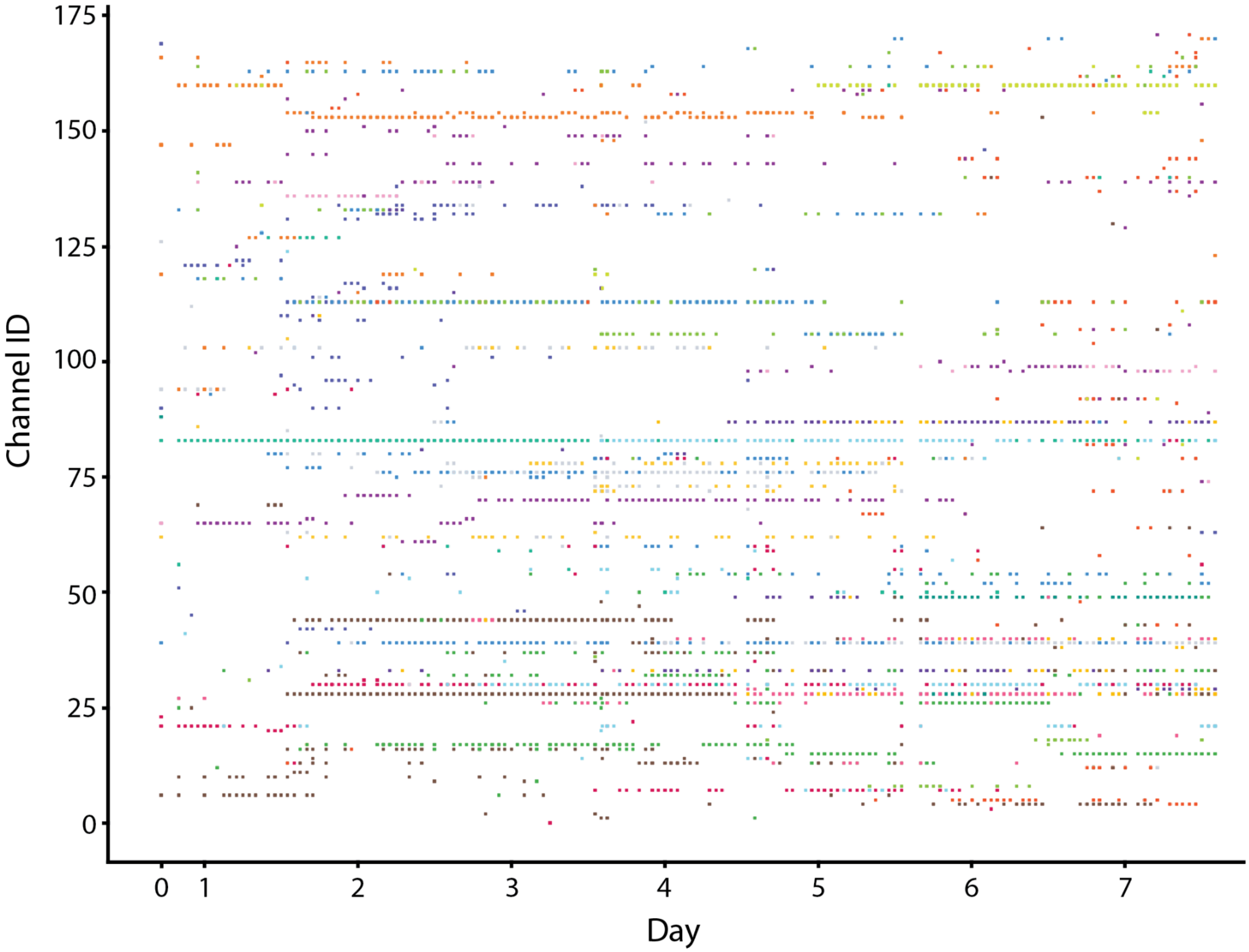


**Figure S6:** **Waveform cluster channel location to recording.**

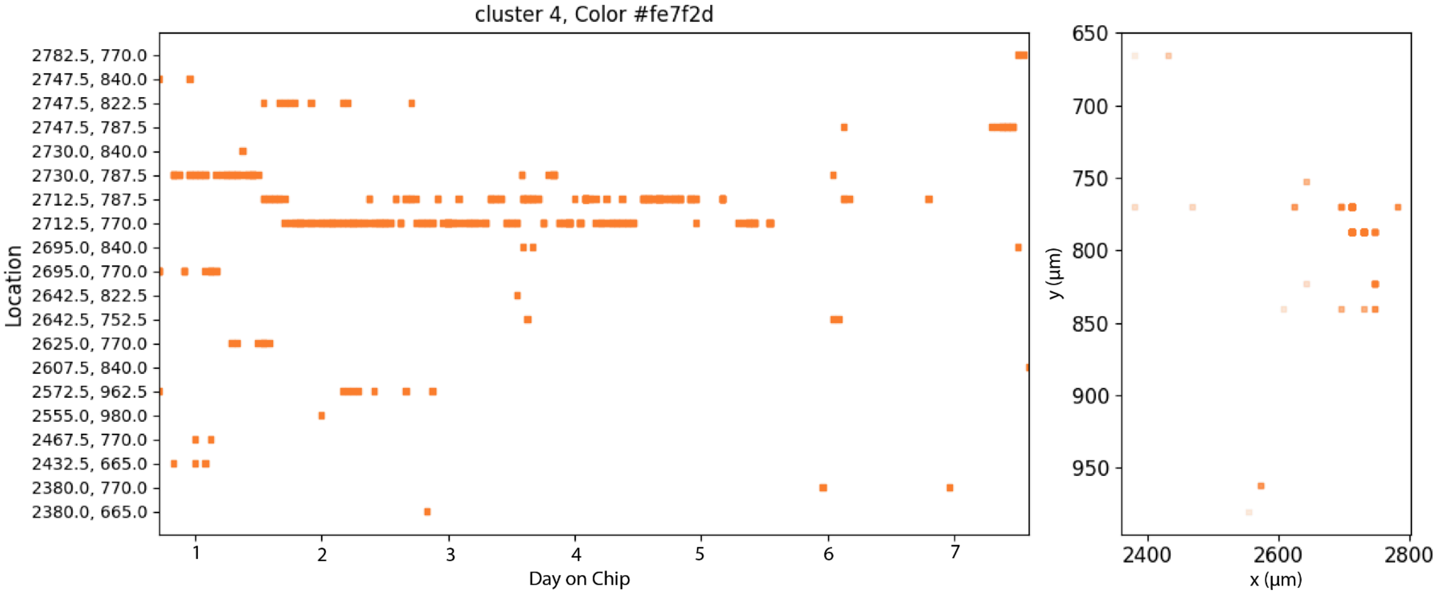


**Figure S7:** **Channel location to recording for Cluster 4**

#### Optogenetics Modulation of Epileptiform Activity

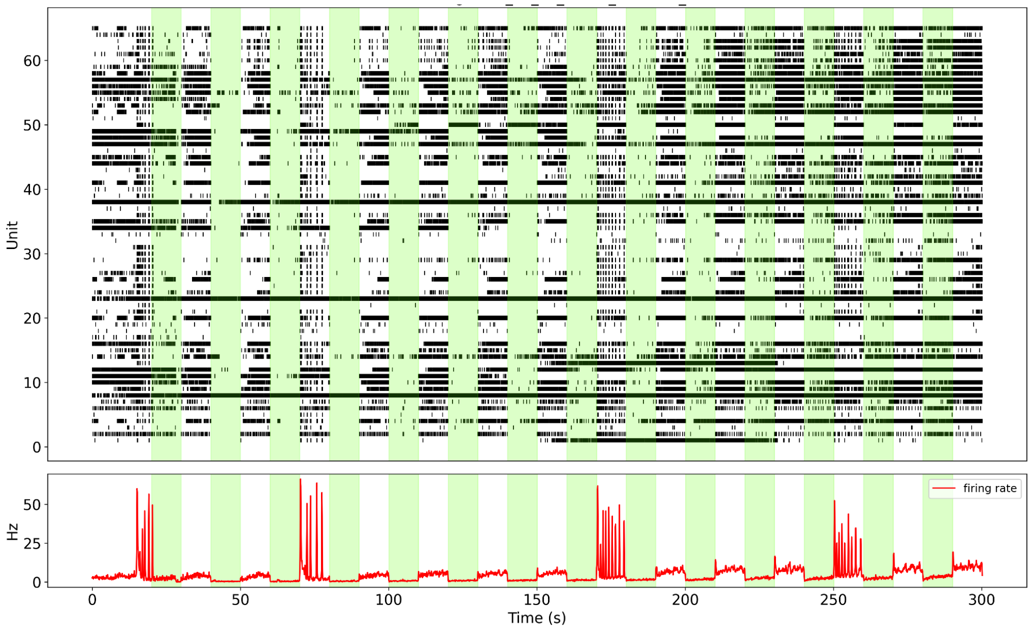


**Figure S8: Spike raster and population firing rate showing neuronal activity modulation under optogenetics illumination.**
